# Annotating the genome at single-nucleotide resolution with DNA foundation models

**DOI:** 10.1101/2024.03.14.584712

**Authors:** Bernardo P. de Almeida, Hugo Dalla-Torre, Guillaume Richard, Christopher Blum, Lorenz Hexemer, Maxence Gélard, Javier Mendoza-Revilla, Ziqi Tang, Frederikke I. Marin, David M. Emms, Priyanka Pandey, Stefan Laurent, Marie Lopez, Alexandre Laterre, Maren Lang, Uğur Şahin, Karim Beguir, Thomas Pierrot

## Abstract

Genome annotation models that directly analyze DNA sequences are indispensable for modern biological research, enabling rapid and accurate identification of genes and other functional elements. This capability is paramount as the volume of sequenced genomes rapidly expands, making the need for efficient and accurate annotation methods increasingly critical, particularly in the context of genetic variant prediction and in-silico sequence design. Current annotation tools are typically developed for specific element classes and trained from scratch using supervised learning on datasets that are often limited in size. This approach constrains their performance and ability to generalize to new genomes. Here, we frame the genome annotation problem as instance segmentation and introduce a novel methodology for fine-tuning pre-trained DNA foundation models to segment 14 different genic and regulatory elements at single-nucleotide resolution. We leverage the self-supervised pre-trained model Nucleotide Transformer (NT) to develop a general segmentation model, SegmentNT, capable of processing DNA sequences up to 50kb long. By utilizing pre-trained weights from NT, SegmentNT surpasses the performance of several ablation models and baselines, including convolutional networks with one-hot encoded nucleotide sequences and large models trained from scratch. We demonstrate state-of-the-art performance on gene annotation, splice site and regulatory elements detection throughout the genome. We also leveraged our framework to accommodate two extra DNA foundation models, Enformer and Borzoi, extending the sequence context up to 500kb and enhancing performance on regulatory elements. Finally, we show that a SegmentNT model trained on human genomic elements generalizes to elements of different species, and a multi-species SegmentNT model achieves strong generalization across unseen species. Our approach is readily extensible to additional genomic elements and species. We have made our SegmentNT human and multi-species models, as well as the SegmentEnformer and SegmentBorzoi models, available on our github repository in Jax and HuggingFace space in Pytorch.

## Introduction

Genome annotation models play a pivotal role in modern biological research, providing the tools necessary to identify genes, their exon-intron structure, and other functional elements directly from DNA sequences. This capability is becoming increasingly essential as the volume of sequenced genomes continues to grow exponentially, driven by advancements in sequencing technologies [1]. Accurate and efficient annotation of DNA sequences not only facilitates the understanding of genetic architecture but is also critical for applications such as genetic variant prediction and in-silico sequence design.

Current annotation pipelines, such as BRAKER2 [2] and MAKER2 [3], rely on *de novo* sequence-based predictors using Hidden Markov Models (HMMs), including models like Genscan [4] and AUGUSTUS [5, 6]. Although these models have single-nucleotide resolution, they lack the capacity to fully model biological complexity on their own and thus have limitations on predicting gene isoforms and when applied through whole chromosomes, requiring the integration with experimental data (e.g. RNA-seq) and homology to previously characterized proteins to improve accuracy. In addition, these models are focused on gene elements and cannot annotate other types of genomic elements such as regulatory regions. This narrow focus restricts their performance and reduces their ability to generalize to novel or under-represented genomes.

Alternative methods for identifying regulatory elements like promoters [7, 8], enhancers [9–12] or polyA signals [13–15] are usually developed for each specific element class separately and trained from scratch using supervised learning on datasets that are often limited in size. In addition, many of these tools are trained on curated datasets with different distribution from the use-case scenario, significantly hindering their performance when applied to actual genomes. As a result, there is a pressing need for more versatile approaches that can overcome these challenges, generalize to all relevant genomic element types, and adapt to the rapidly expanding landscape of genomic data. A model that can learn sequence dependencies directly from the DNA and efficiently annotate complete genomes with high accuracy would not only streamline genome annotation processes but also deepen our understanding of the genomic code.

The intersection of genomics research and deep learning methods is profoundly changing our ability to understand the information encoded in the human genome [16, 17]. The abundance of available sequencing and omics data have recently led to the development of DNA foundation models — flexible AI models pre-trained on broad data sets that can be applied to a variety of genomic tasks. These include models trained in a supervised manner on thousands of experimental data (e.g. Enformer [18] and Borzoi [19]) or trained in a self-supervised setting across various unlabeled genome sequences [20–26]. This last approach in particular is very promising for genomics given the ability of such foundation models to be trained on unlabeled data (e.g. raw genomes or experimental sequencing data), creating general-purpose representations capable of solving a multitude of downstream tasks, similarly to what has been observed in other fields such as natural language processing and computer vision [27–31].

In this work, we explore the use of DNA foundation models and their learned representations to develop versatile models capable of annotating the location of several types of elements in the genome at single-nucleotide resolution. Given the similarities between localizing elements at nucleotide resolution in a DNA sequence and localizing objects in images at pixel resolution, usually referred to as segmentation task [32–34], we framed the genome annotation problem as instance segmentation and adopted a segmentation architecture that proved useful in that field. More specifically, we built a DNA segmentation model, the Segment-Nucleotide Transformer (SegmentNT), that combines the pre-trained DNA foundation model Nucleotide Transformer (NT) [22] and a 1D U-Net [32] architecture, and trained it to predict the location of 14 types of human regulatory and gene elements in input sequences up to 30kb at single-nucleotide resolution. We show that SegmentNT achieves high performance on gene annotation, splice site and regulatory elements detection throughout the genome and generalizes to input sequences up to 50kb. Our framework is compatible with different DNA encoders, as we demonstrate by integrating two additional DNA foundation models, Enformer and Borzoi, that allowed to extend the sequence context to 500kb, enhancing performance on regulatory element detection. We further fine-tuned our best SegmentNT-30kb model on multiple species and show improved generalization to unseen animal and plant species. Our framework is general and readily extensible to additional DNA foundation models, genomic elements and species genomes.

## Results

### SegmentNT: fine-tuning Nucleotide Transformer for segmentation of DNA sequences at nucleotide resolution

We developed a new model called SegmentNT to annotate the location of several types of genomic elements in a sequence at single-nucleotide resolution. Following segmentation principles, we framed this problem as instance segmentation, computing a binary mask over nucleotides for each type of element. SegmentNT combines the pre-trained DNA foundation model Nucleotide Transformer (NT) [22] and a segmentation head to detect elements at different scales (Fig. 1a). As segmentation head we make use of a 1D U-Net architecture that downscales and upscales the foundation model embeddings of the input DNA sequence (Fig. 1b; see also Linder et al.[19] for a recent use-case of U-Net in genomics). This architecture is trained end-to-end on a dataset of genomic annotations to minimize a focal loss objective [33] to deal with element scarcity in the dataset (see Methods).

**Figure 1.**
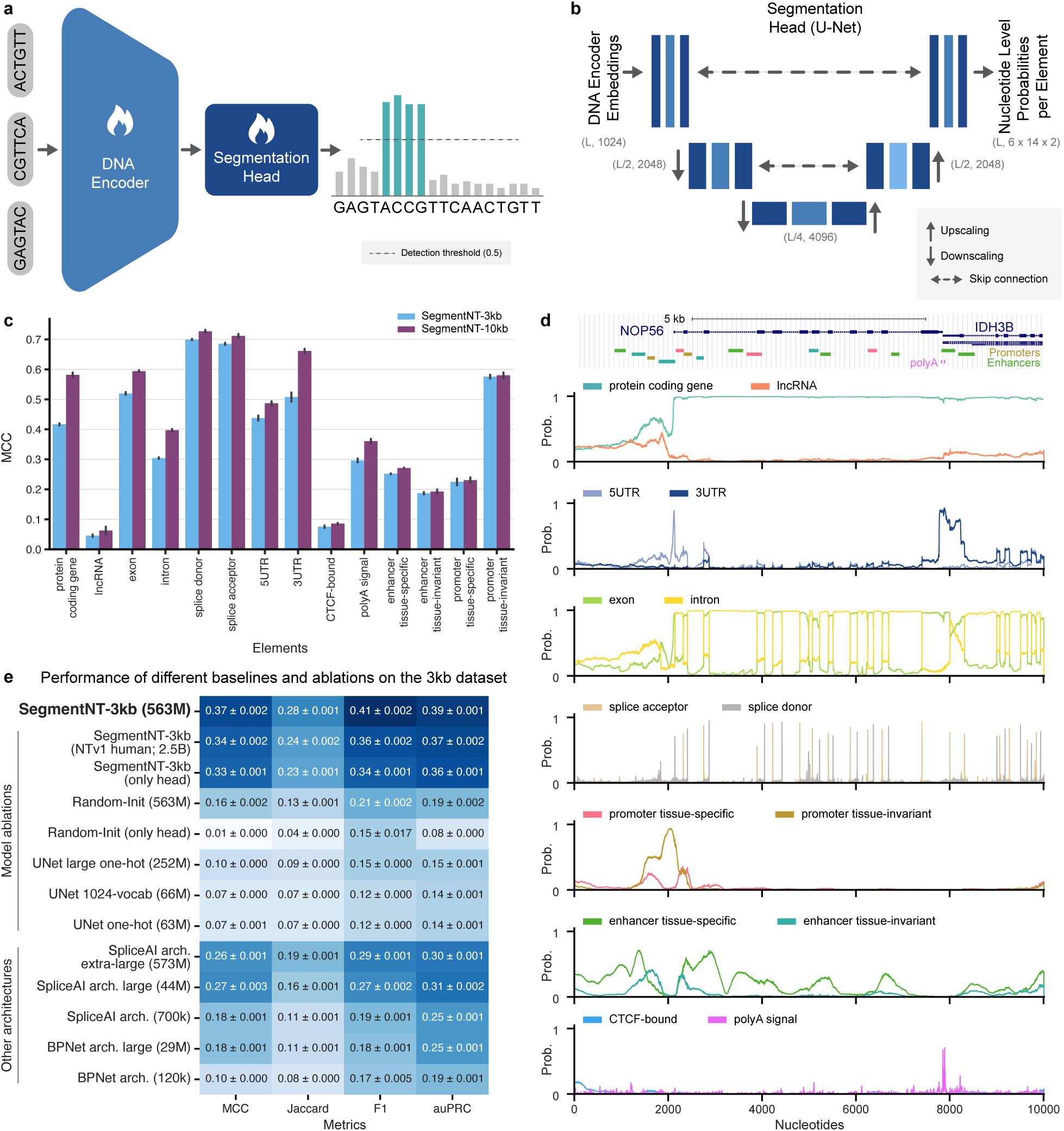
SegmentNT localizes genomic elements at nucleotide resolution. **a)** The SegmentNT neural network architecture consists of a pre-trained DNA encoder (here Nucleotide Transformer (NT) [22]) and a segmentation head (here a U-Net). The output are probabilities for each genomic element at nucleotide resolution. **b)** As segmentation head we use a 1D U-Net architecture with 2 downsampling and 2 upsampling convolutional blocks with matched U-Net connections. We added the dimensions of each layer. **c)** Performance of SegmentNT trained on 3kb and 10kb sequences on 14 types of genomic elements. We used as metric the Matthews correlation coefficient (MCC). Data are presented as mean MCC values +/- 95% confidence interval from 10 different samplings of the test set. **d)** Representative example of annotations and predicted probabilities of the 14 types of genomic elements at the *NOP56*/*IDH3B* gene locus located in the test set. Gene isoforms with respective exons and introns, as well as promoter and enhancer regulatory elements are shown. **e)** Comparison of performance between SegmentNT and different model ablations and architectures. The metrics used are the MCC, Jaccard index, F1 score and the area under the precision-recall curve (auPRC). Performance for each metric is presented as mean metric values +/- standard deviation across the 14 different element types.

To train SegmentNT we curated a dataset of annotations at nucleotide-level precision for 14 types of genomic elements in the human genome derived from GENCODE [35] and ENCODE [36], including gene elements (protein-coding genes, lncRNAs, 5’UTR, 3’UTR, exon, intron, splice acceptor and donor sites) and regulatory elements (polyA signal, tissue-invariant and tissue-specific promoters and enhancers, and CTCF-bound sites) (Supplementary Fig. 1; see Methods). Since these element annotations can overlap, SegmentNT predicts separately the probability of belonging to each of the genomic elements at nucleotide level. For example, in different gene transcript isoforms the same DNA region can be considered an exon or an intron, enhancers can also be found in gene regions, and polyA signals are usually in the gene’s 3’UTRs. In addition, here we used the canonical definition of exons as any part of a gene that can be present in the final mature RNA after introns have been removed by RNA splicing, thus also overlapping with 5’ and 3’UTRs. This allows the prediction of every genomic element independent of the other predictions. The annotation of all promoter and enhancer regions in the human genome was derived from the latest registry of candidate cis-regulatory elements by ENCODE [37]. It contains 790k enhancers and 34k promoters grouped by their activity in different tissues.

We first trained a model to segment these distinct 14 genomic elements in input DNA sequences of 3kb (SegmentNT-3kb). This model was further fine-tuned on 10kb input sequences (SegmentNT-10kb) to extend its input length. This was achieved by initializing SegmentNT-10kb from the best checkpoint of the SegmentNT-3kb model for a more efficient training and length-adaptation. For a given input sequence, these models make 42, 000 and 140, 000 predictions, respectively, each being the probability of a given nucleotide to belong to a genomic element type. For binary classification metrics we used as threshold 0.5 to annotate nucleotides as belonging to each element type. Model training, validation and performance evaluation were performed on different sets of chromosomes from the human genome, accounting for homologous sequences to ensure no data leakage and that the test set provides a robust evaluation of model performance. The models were evaluated on ten different samplings of the test set and the performance was assessed using each nucleotide as a separate prediction and with various metrics: Matthews correlation coefficient (MCC), area under the precision-recall curve (auPRC), Jaccard similarity and the F1-score (see Methods; Supplementary Fig. 2). SegmentNT-3kb demonstrated high accuracy in localizing the different elements to nucleotide precision, showing a Matthews correlation coefficient (MCC) on the test set above 0.5 for exons, splice sites, 3’UTRs and tissue-invariant promoter regions (Fig. 1c). LncRNA and CTCF-binding sites were the most difficult elements to predict, with test MCC values below 0.1. We observed superior performance of the model in sequences of 10kb (average MCC of 0.42) compared with 3kb (0.37), in particular for protein-coding genes, 3’UTRs, exons and introns, suggesting that these elements depend on longer sequence contexts (Fig. 1c). These results were consistent across the different metrics (Supplementary Fig. 2).

To further evaluate predictive performance, we inspected regions of the held-out test chromosomes. Evaluating SegmentNT-10kb on a 10kb window that covers the gene *NOP56* on the positive strand and the end of the gene *IDH3B* on the negative strand shows that it accurately predicts the different genic elements of each gene (Fig. 1d). SegmentNT correctly predicts both genes as protein-coding, their 5’UTR and 3’UTR positions, their splice sites and exon-intron structure, and also the polyA signals. In addition, SegmentNT captures the promoter region of *NOP56*, both the tissue-specific and tissue-invariant ones. This region also contains multiple enhancers and some of those are correctly predicted by the model. Still, although our global performance metric for enhancers is good (MCC of 0.27 for tissue-specific and 0.19 for tissue-invariant for SegmentNT-10kb), we observe that enhancer predictions are more noisy. This could be related to their higher sequence complexity and diversity, and we expect that grouping them by cell type-specific activity should further improve model performance (see Discussion). Additional comparisons between SegmentNT models and competitive approaches for gene annotation, splice site prediction and detection of regulatory elements are detailed in later sections.

### Using a pre-trained DNA encoder is essential for efficient training and to achieve superior performance

We next evaluated our model design choices and the importance of using the NT pre-trained foundation model as a DNA encoder. We present all model baseline and ablation architectures and results in Supplementary Tables 1, 2 and 3, and statistical tests in Supplementary Fig. 3. We first compared the performance of SegmentNT with different model ablations, using 3kb input sequences for a simpler comparison (see Methods). We removed the NT DNA encoder and trained two 1D U-Net architectures that take one-hot encoded DNA sequences directly as input instead of the NT embeddings: one with the same 63M parameters of the head of SegmentNT and a larger one with an additional downsampling/upsampling block featuring a total of 252M parameters. We tested an additional one that first expands the raw DNA sequence input to the same vocabulary dimension of the NT embeddings (1,024) before passing it through the same U-Net architecture as SegmentNT. These three U-Net architectures demonstrated substantially reduced performance across all elements, with an average MCC of 0.07 (66M), 0.11 (250M) and 0.07 (1024-vocab) compared with 0.37 for SegmentNT-3kb, demonstrating the value of using a DNA encoder (Fig. 1e).

To test the benefit of pretraining the NT foundation model, we trained a model version with the same architecture as SegmentNT but using a randomly initialized NT DNA encoder model, rather than the pre-trained one. Our ablation study shows that on this task, self-supervised pre-training on genomes allows our SegmentNT model to converge 7 times faster to an asymptotic performance twice greater across all 14 genomic elements: average MCC 0.37 compared with 0.16 for the version with a random initialized NT (Fig. 1e). Additional ablation analyses showed that (1) fine-tuning both the NT DNA encoder and the U-Net head achieves higher performance (Fig. 1e), (2) SegmentNT using the NT pre-trained on multispecies genomes is superior to one using a NT model that was pre-trained on thousands of human genomes (NTv1 human [22]; Fig. 1e).

Finally, we compared SegmentNT with two other popular, small CNN architectures in genomics developed for nucleotide-resolution tasks that do not use pre-trained models: BPNet [38] and SpliceAI [39] that proved successful in modeling transcription-factor binding and splicing, respectively. We used their original archi-tectures (BPNet 120k and SpliceAI 700k parameters) and scaled versions where we increase the embedding dimensions across all their layers (see Methods). Despite the small scale, the initial SpliceAI architecture (average MCC 0.18) achieved improved performance over both BPNet (0.10), U-Net (0.07) and SegmentNT with a randomly initialized NT encoder model (0.16, Fig. 1e). We could further increase its performance to an average MCC of 0.27 by scaling the model embeddings, showing its effective architecture among supervised CNN models, however still far from our SegmentNT approach (0.37). We note that these CNN architectures could still be further optimized to this multi-element segmentation task, but it is out-of-scope of this work. Overall, the superior performance of SegmentNT over these different model baselines demonstrates the value of DNA foundation models for solving challenging tasks in genomics such as localizing different types of genomic elements at a single-nucleotide resolution.

### SegmentNT generalizes to sequences up to 50kb

We next investigated how to extend the sequence context length of SegmentNT, motivated by the improved results observed for SegmentNT-10kb over SegmentNT-3kb and the long-range interactions prevalent in the humane genome (Fig. 1c). Given that NT uses rotary positional embeddings (RoPE; [40]) that were set to support sequences up to 12kb during its pre-training, using NT directly on sequences longer than 12kb, whether for fine-tuning or inference, would yield poor performance due to the periodic nature of RoPE encoding. To address this problem, we explored recent approaches that have been proposed for extending contexts of RoPE models by converting the problem of length extrapolation into one of “interpolation”. Specifically, we employ a context-length extension method first formally described in [41], where the frequency used in RoPE embeddings is re-scaled to account for longer sequences (see also [42, 43]). This approach can be used for extending the context length of SegmentNT during the fine-tuning on sequences longer than 12kb but also for performing inference with SegmentNT models on sequences longer than the ones seen during training. We investigated both scenarios below.

We implemented context-length extension in NT and trained two additional SegmentNT models that segment the 14 genomic elements in DNA sequences of 20kb (SegmentNT-20kb) and 30kb (SegmentNT-30kb) (see Methods). Evaluation on the same test chromosomes showed consistent improvements in performance with increased sequence length, in particular for the segmentation of protein-coding genes, 3’UTRs, exons and introns (Fig. 2a, Supplementary Fig. 3, Supplementary Tables 2 and 3). The model with the best performance across all elements was SegmentNT-30kb with an average MCC of 0.45 (Fig. 2b).

**Figure 2.**
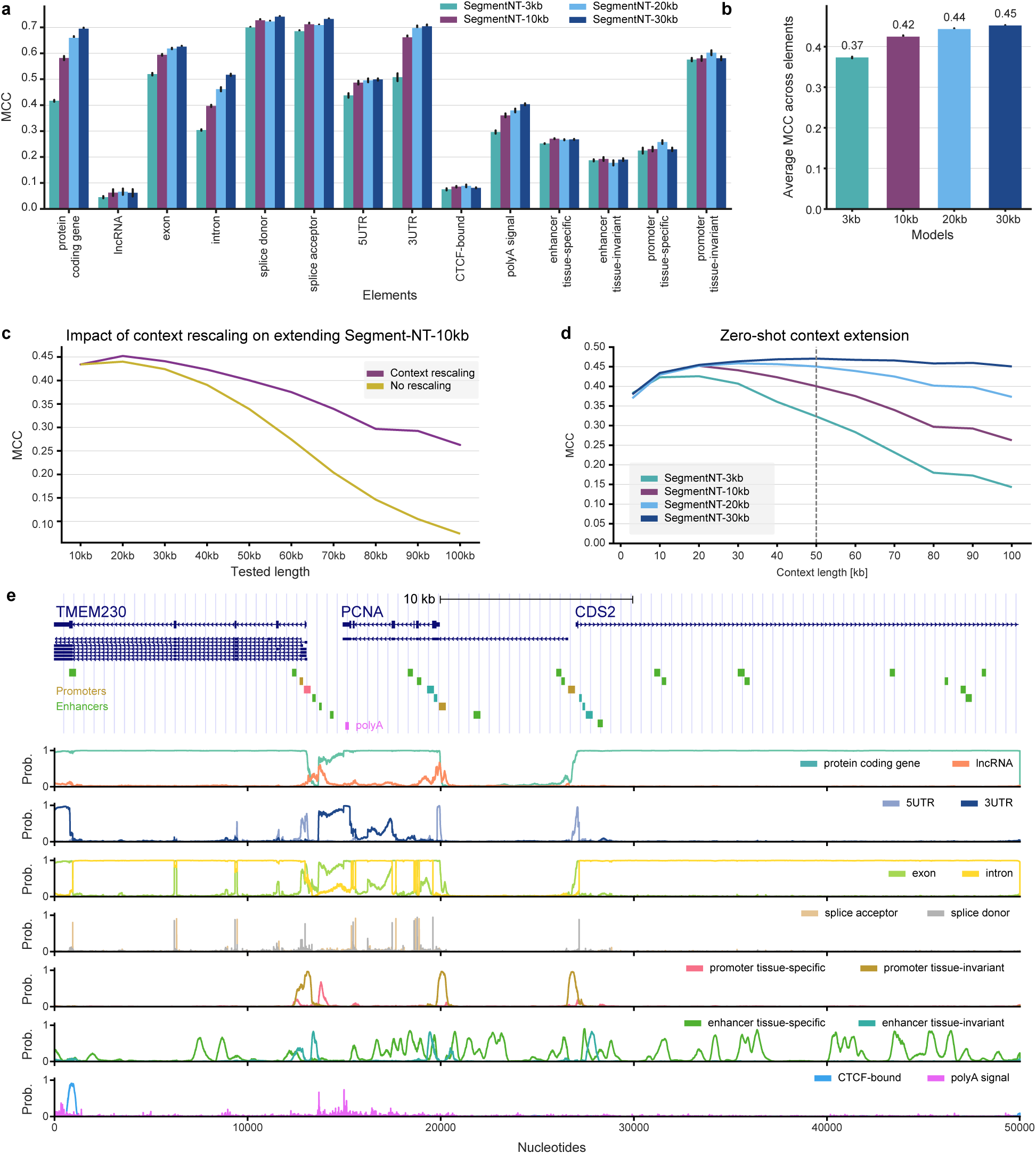
Adaptation and zero-shot generalization of SegmentNT across multiple sequence lengths. **a)** Performance of SegmentNT trained on 3kb, 10kb, 20kb and 30kb sequences on 14 types of genomic elements. We used as metric the MCC. Data are presented as mean MCC values +/- 95% confidence interval from 10 different samplings of the test set. **b)** Average MCC performance of the different models across the 14 elements. Data are presented as mean MCC values +/- 95% confidence interval from the 14 elements. **c)** Context-length extension through Rotary Position Embedding (RoPE) rescaling allows to improve performance of SegmentNT-10kb on up to 100kb sequences. Average MCC performance across the 14 elements for the SegmentNT-10kb model with and without context-length rescaling. **d)** Long-range models improve generalization on longer contexts while maintaining performance on shorter contexts. Average MCC performance across the 14 elements for the different SegmentNT models per input sequence length. **e)** Representative example of annotations and predicted probabilities of the 14 types of genomic elements for a 50kb region at the *TMEM230*/*PCNA*/*CDS2* gene locus located in the test set. Gene isoforms with respective exons and introns, as well as promoter and enhancer regulatory elements are shown.

Since it is computationally expensive to fine-tune SegmentNT on even longer sequence lengths, we tested if we could leverage context-length extension to evaluate a model pre-trained on a given length on longer sequences. We tested this approach on the SegmentNT-10kb model and evaluated it with or without context-length extension on the prediction of sequences up to 100kb from the same test chromosomes (Fig. 2c, Supplementary Fig. 4). Context-length extension substantially improved the performance of the model on longer sequences, in particular on 100kb where the original model showed very poor performance (average MCC of 0.26 vs 0.07, respectively).

This motivated us to more systematically test how far our different SegmentNT models could be extended. To address that, we evaluated the performance of all trained SegmentNT models (3kb, 10kb, 20kb and 30kb) on input sequence lengths between 3 and 100kb using context length extension interpolation when needed. When averaging the performance across 14 elements, this revealed that the model trained on the longest context length (SegmentNT-30kb) achieved the best results when evaluated in all context lengths, including shorter sequences (Fig. 2d). We observed top performance for 50kb input sequences (average MCC of 0.47) and a drop in performance for sequences longer than 50kb, although SegmentNT-30kb still has good performance on sequences of 100kb (0.45; Fig. 2d). These results highlight the flexibility of SegmentNT and how it can be applied to sequences of different lengths. We note that the SegmentNT-30kb model when segmenting the 14 genomic elements in an 50kb input sequence makes 700, 000 predictions at once (14 x 50, 000), thus providing a very rich segmentation output. See an example of the SegmentNT-30kb predictions for a 50kb locus in the test set with three overlapping genes (Fig. 2e).

Finally, we have used our best 30kb longer-range model to investigate in more detail the performance gaps of SegmentNT and if the mispredictions are coming from edge effects of annotations, in particular for regulatory elements where hard boundaries are more arbitrary, or spurious predictions in random locations. In order to do this, per type of element, we have calculated the enrichment of mispredictions at regions edges, inside regions, or in random locations outside labeled regions. We observed for all elements a strong enrichment of mispredictions at edge nucleotides but also at nucleotides inside the labeled regions. For all regulatory element classes, the enrichment of mispredictions inside elements was higher than at the edges, suggesting that the performance gaps come from worse predictions in some whole regions rather than poorly predicted edge effects across regions (Supplementary Table 4).

### Using different foundation models as DNA encoders to extend segmentation to 500kb sequences

Although SegmentNT demonstrates broad generalizability across various types of genomic elements, leveraging sequence representations from the NT pre-trained DNA encoder, its capacity to process longer sequences is constrained by the limitations of the DNA encoder itself (50kb, as demonstrated above; Fig. 2d). Additionally, the performance on certain types of elements studied here could potentially be improved by utilizing DNA encoders that better capture their specific sequence features. Therefore, we investigated alternative models as DNA encoders within our framework, aiming to both extend the input sequence length and assess their generalization across diverse genomic elements.

We compared SegmentNT’s performance when using NT, Enformer [18] and Borzoi [19] as DNA encoders (Fig. 3a). In contrast to NT, Enformer and Borzoi are long-range models pre-trained in a supervised manner to predict thousands of epigenetic and gene expression tracks in various mouse and human cell types, and thus might have learned better representations of regulatory elements for instance. Both models integrate CNNs with a transformer architecture, processing long sequences as input (Enformer: 196kb, Borzoi: 524kb) but predicting at lower resolution (Enformer: 128bp, Borzoi: 32bp). For a systematic comparison with SegmentNT, we have combined the pre-trained representations from their last layer, before the prediction heads, to our U-Net segmentation head and fine-tuned the whole network on our segmentation dataset using either the same 30kb input sequences or the model’s original input length, 196kb for Enformer and 524kb for Borzoi (see Methods, Supplementary Fig. 3 and Supplementary Tables 1, 2 and 3). We named these new architectures

**Figure 3.**
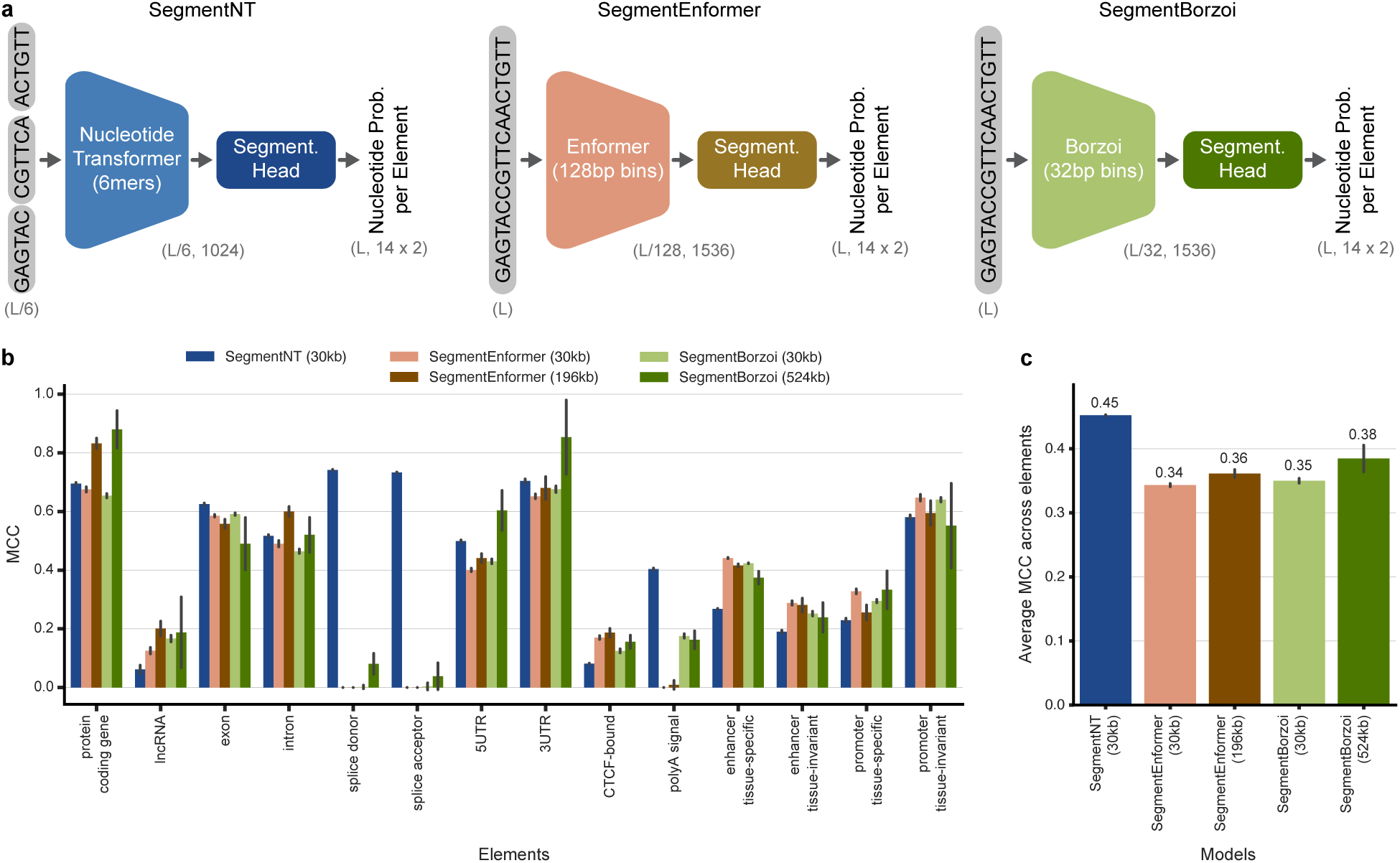
SegmentNT localizes genomic elements at nucleotide resolution. **a)** Cartoons displaying how the different DNA encoders are combined with the U-Net segmentation head to create SegmentNT, SegmentEnformer and SegmentBorzoi models. The model input and output dimensions are shown within parentheses. L: input sequence length. Since each output token in Enformer and Borzoi represents 128bp and 32bp bins, respectively, the segmentation head needs to predict the respective number of probabilities per element per nucleotide, i.e. 14 x 2 x 128 for Enformer and 14 x 2 x 32 for Borzoi. **b)** Performance of the different models on the 14 types of genomic elements. We used as metric the MCC. Data are presented as mean MCC values +/- 95% confidence interval from 10 different samplings of the test set. **c)** Average MCC performance of the different models across the 14 elements. Data are presented as mean MCC values +/- 95% confidence interval from the 14 elements.

#### SegmentEnformer and SegmentBorzoi models for simplicity

We first compared the performance of the three models using 30kb input sequences (Fig. 3b). On average, SegmentNT (average MCC of 0.45) achieved superior performance than SegmentEnformer (0.34) and Segment-Borzoi (0.35; Fig. 3c). However, their performances varied across the different genomic elements. SegmentNT outperformed the others in identifying gene elements, including protein-coding genes, 5’UTR, 3’UTR, exons, introns, splice acceptor and donor sites, and polyA signals. Notably, for splice site and polyA signal prediction — tasks that require high resolution to accurately annotate single- or few-nucleotide elements —SegmentEnformer and SegmentBorzoi performed poorly, likely due to their pre-segmentation resolution limits of 128bp and 32bp bins, respectively. In contrast, SegmentEnformer and SegmentBorzoi showed improved segmentation of lncRNAs, CTCF-bound sites, and promoter and enhancer regulatory elements (Fig. 3b). This improved performance reflects the supervised pre-training of Enformer and Borzoi on epigenomic and transcriptomic profiles, which enhanced their ability to represent these specific sequence elements, supporting our hypothesis that different DNA encoders capture distinct features and can be more suitable for certain types of elements.

We next analysed the performance of SegmentEnformer and SegmentBorzoi with extended input sequences of 196kb and 524kb, respectively. Both models showed an overall performance improvement compared to their 30kb versions (Fig. 3b,c), largely driven by enhanced accuracy in identifying protein-coding genes, lncRNAs, and introns — longer elements that previously also benefited from increased sequence length (Fig. 2a). SegmentBorzoi demonstrated additional gains in both UTR regions, likely due to its pre-training with RNA-seq data that allowed it to learn comprehensive features of gene structure and all types of transcribed regions [19]; unlike Enformer, which was pre-trained using only CAGE expression data that primarily captures the start of gene transcription [18]. As anticipated, performance on regulatory elements did not improve significantly, as these elements do not typically require long-distance interactions. Despite those improvements, the average performance of both models across genomic elements was still significantly lower than SegmentNT (Fig. 3c). Overall, our findings emphasize that specific DNA encoders can be leveraged to achieve superior performance on particular genomic elements, while SegmentNT remains the best default approach due to its consistent generalization across all element types.

#### Comparison with established gene annotation tools

After having explored the different aspects of the SegmentNT architecture, we investigated the performance of our best SegmentNT-30kb model in established gene annotation tasks. Here, we compared our model with the established HMM gene finder AUGUSTUS [5, 6], state-of-the-art among sequence-based gene finders [3]. We first evaluated the two models in the gene annotation task presented at the recent BEND benchmark [44], analysing only sequences in the SegmentNT test chromosomes. Since our model predicts all gene isoforms but this task considers only the main ioform of each gene, we have adapted it to create a version with only genes that contain a single isoform (Fig. 4a,b, Supplementary Fig. 5a) and a version that contains all genes and respective isoforms, allowing multiple labels per nucleotide (Fig. 4c, Supplementary Fig. 5b,c; see Methods). We measured their performance using the standard F1-score and the MCC metric, using 0.5 as the probability threshold.

**Figure 4.**
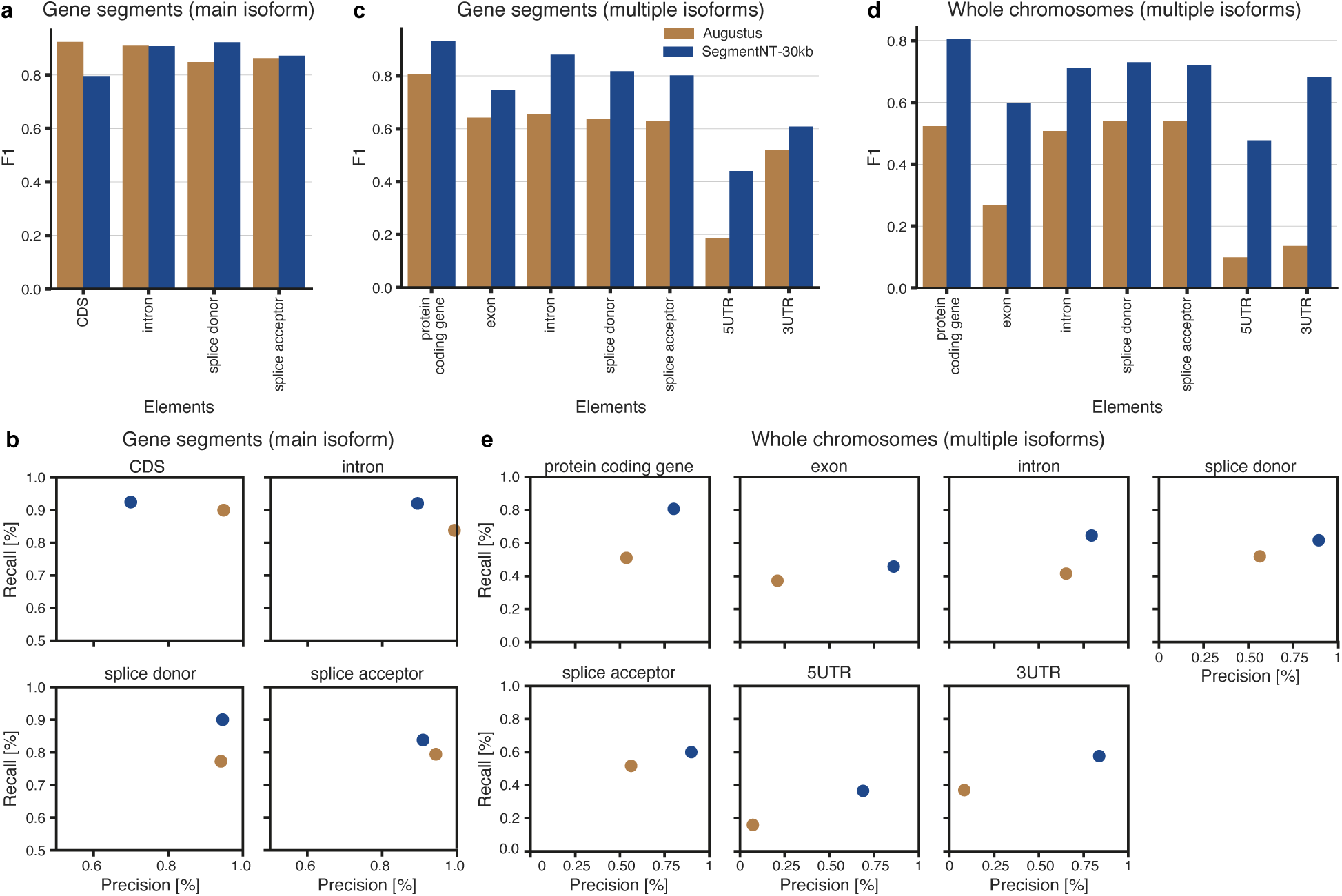
Comparison with AUGUSTUS for gene prediction. **a,c,d)** Accuracy of SegmentNT-30kb and AUGUSTUS for the different gene elements in different datasets: 30kb segments with genes using only their main isoform **(a)**, 30kb segments with genes considering all annotated gene isoforms **(c)** and whole segmentation test chromosomes considering all annotated gene isoforms **(d)**. The metric used was the F1-score. **b)** Precision and recall for SegmentNT-30kb and AUGUSTUS for the different gene elements in the dataset of 30kb segments with genes using only their main isoform. **e)** Precision and recall for SegmentNT-30kb and AUGUSTUS for the different gene elements in the whole segmentation test chromosomes, considering all annotated gene isoforms.

Our results demonstrated that SegmentNT-30kb is competitive with AUGUSTUS for the easier setting of segmenting the main isoform of various genes, achieving improved performance for splice donor sites, similar performance for introns and splice acceptor sites, and worse performance for CDS regions (Fig. 4a,b, Supplementary Fig. 5a). The latter is due to lower precision despite having similar recall (Fig. 4b). When considering all confident gene isoforms, SegmentNT-30kb outperforms the HMM model in all gene elements in both metrics (Fig. 4c, Supplementary Fig. 5b,c). In all cases, SegmentNT-30kb has higher recall, most of the times accompanied by higher precision.

Finally, we compared both models along the test set settings of SegmentNT, where the model needs to segment the gene regions along the whole test set chromosomes, where most regions do not contain genes, together with their respective isoforms (Fig. 4d,e, Supplementary Fig. 5d). Also here, SegmentNT-30kb outperformed AUGUSTUS and achieved superior performance across all gene elements by a large margin, with both higher recall and precision. This result agrees with the limitations of this type of tools when scanned through the whole genome, in contrast to their very high accuracy when segmenting a genic region, and highlights the superior performance of SegmentNT on the complex task of whole genome segmentation.

#### SegmentNT accurately predicts splice sites across the genome

One of the main nucleotide-level tasks in genomics that has been tackled by previous models is splice site detection, where SpliceAI [39] and Pangolin [45] are considered state-of-the-art. We first compared SegmentNT-30kb with the specialized SpliceAI-10kb and Pangolin models on detecting splice donor and acceptor nucleotides on a gene from our test set (*EBF4*; Fig. 5a). SegmentNT correctly predicts all exons and introns in addition to all splice sites, including the ones of the alternative exon at the gene start. When comparing the different models we observe that SpliceAI and Pangolin predict all existent splice sites but overpredict additional sites (see red stars in Fig. 5a).

**Figure 5.**
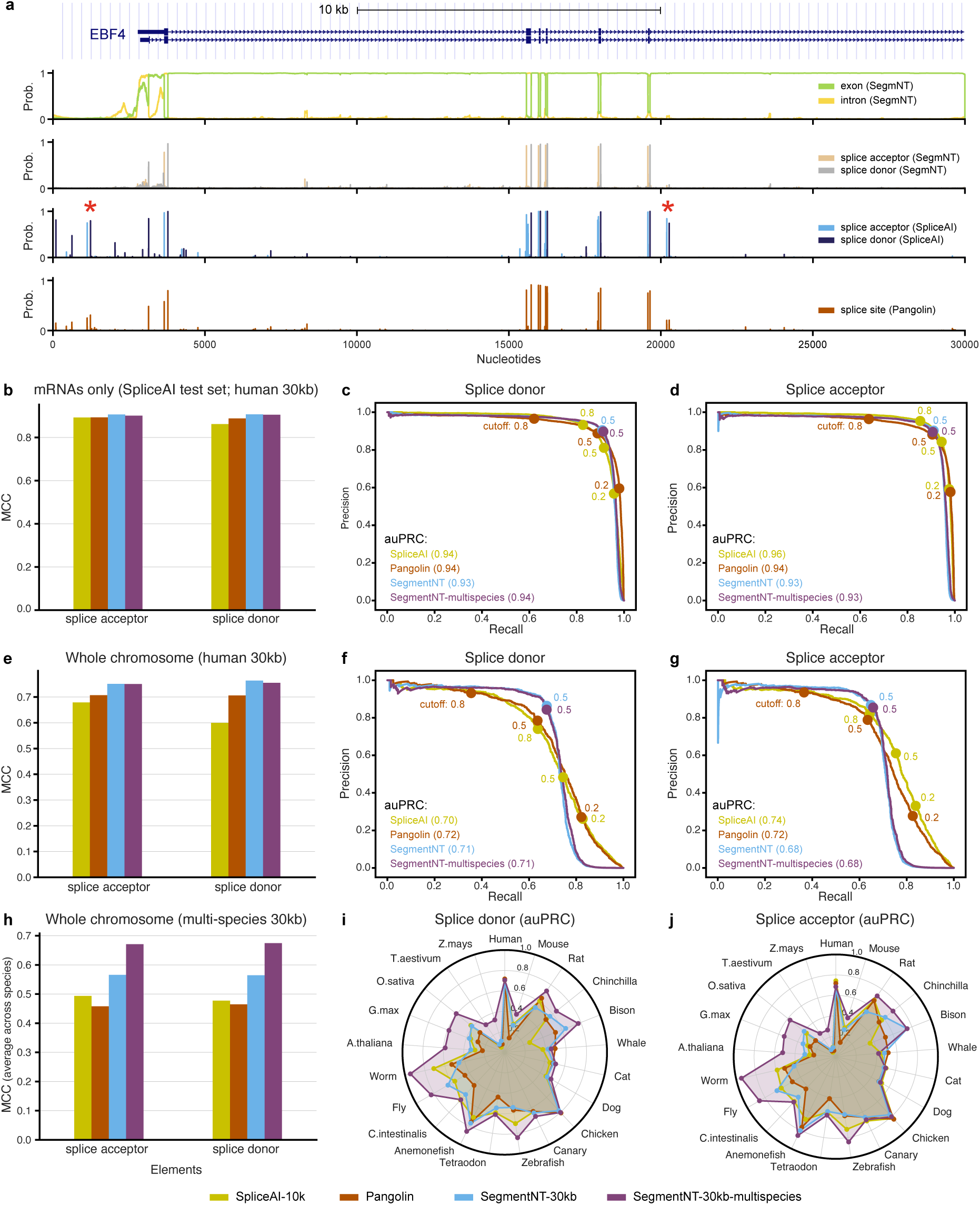
SegmentNT achieves state-of-the performance on splice site prediction. **a)** Representative example of gene annotations and predicted probabilities for splicing elements by SegmentNT-30kb, SpliceAI and Pangolin at the *EBF4* gene locus located in the test set. Gene isoforms with respective exons and introns are shown together with splice site predictions for the different models. SpliceAI/Pangolin mispredicted regions are highlighted with stars. **b,e,h)** Performance of the SpliceAI, Pangolin and SegmentNT-30kb and -multispecies models for splice acceptor and donor detection. We show MCC values on **(a)** the SpliceAI’s mRNA-based test set and the **(b)** human and **(c)** multispecies SegmentNT’s whole genome test sets. The performance in the multispecies dataset is based on the average across 20 species. **c,d,f,g)** Precision-recall curves for splice **(c,f)** donor and **(d,g)** acceptor sites on **(c,d)** the SpliceAI’s mRNA-based test set and the **(f,g)** human SegmentNT’s whole genome test sets. auPRC values are shown along with different thresholds for each model. **i,j)** Radar plots depicting the predictive performance of the four models for splice **(i)** donor and **( j)** acceptor sites. The metric used in the auPRC.

For a comprehensive comparison covering different use-cases, we evaluated each model in both SpliceAI/Pangolin’s test set composed of only mRNA sequences and adapted to 30kb windows, and in our whole chromosome test set, using only genes in the positive strand to match the training scenarios of SpliceAI and Pangolin (see Methods). We note that Pangolin has a different training scheme and does not differentiate between acceptor and donor sites, predicting a single splice site label that we converted into acceptor and donor predictions for a more direct comparison. We evaluated the models’ performance using the standard splicing metrics auPRC, precision-recall curves and top-k accuracy [39], but also MCC as a classification metric using the recommended threshold of 0.5 for all models (Fig. 5, Supplementary Fig. 6).

SegmentNT-30kb achieved comparable performance to SpliceAI and Pangolin on their test set: PR-AUC for donor sites of 0.93 vs 0.94 and 0.94, and for acceptor sites of 0.93 vs 0.96 and 0.94, respectively (Fig. 5b-d, Supplementary Fig. 6a). On SegmentNT’s whole genome test set, our model showed higher classification performance as measured by MCC on both donor and acceptor sites (Fig. 5e). When considering the ranking-based metrics, SegmentNT-30kb had similar auPRC values as SpliceAI and Pangolin for splice donors but lower values for splice acceptor sites (Fig. 5f,g); the top-k accuracy was similar for all models for splice acceptor but was higher for SegmentNT-30kb for splice donor sites (Supplementary Fig. 6b). Despite this high performance overall, we observed lower performance for SegmentNT and Pangolin compared with SpliceAI on non-coding RNA splices, suggesting that the high splicing detection performance could be driven by some correlative signal from coding sequence (Supplementary Fig. 6c). Overall, SegmentNT detects splice donor and acceptor sites in any given input DNA sequence with high accuracy, being more precise on coding sequences across the genome than current methods.

### Localization of regulatory elements

Regarding the detection of regulatory elements like promoters and enhancers, to our knowledge there are no models that can predict the location of such elements in large input sequences at nucleotide resolution. Most existent models are trained on curated datasets with sequences of the same size of promoters or enhancers, significantly hindering their performance in relevant use-cases such as annotating them in actual genomes. We have compared our best segmentation models (SegmentNT, SegmentEnformer and SegmentBorzoi) with competitive approaches that use model classifiers in a sliding-window setting to derive nucleotide-level predictions (Fig. 6a). Here we used as binary classifiers the DeePromoter for promoter prediction [7] as well as the Nucleotide Transformer models fine-tuned on promoter and enhancer sequences that were shown to be state-of-the-art in a previous benchmark [22] (see Methods). We compared these different approaches for the prediction of promoters and enhancers at nucleotide resolution on 30kb input sequences, combining tissue-invariant and tissue-specific ones into a single class since the benchmarked models are global predictors of promoters and enhancers, respectively.

**Figure 6.**
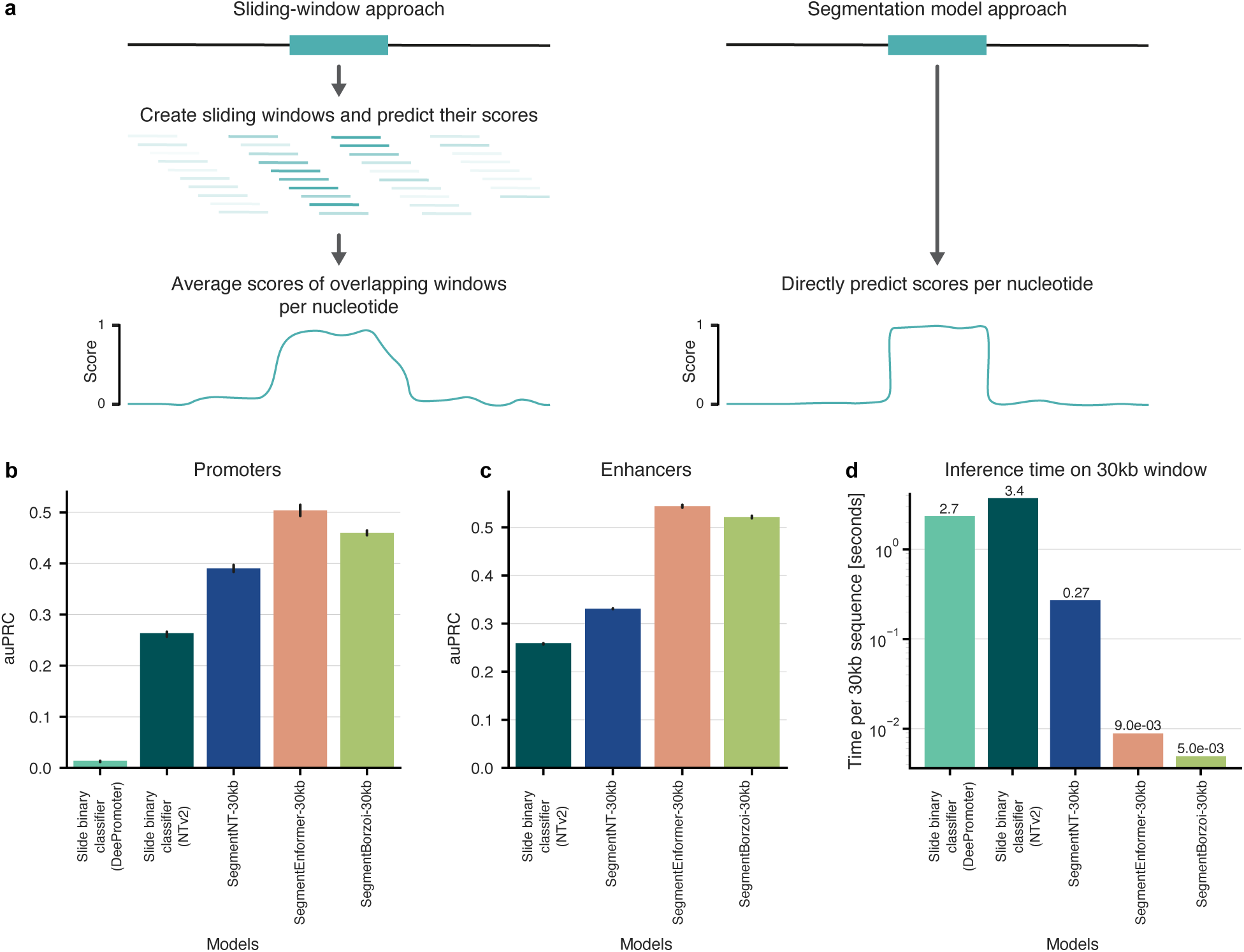
Comparison between segmentation models and alternative sliding-window approaches for promoter and enhancer predictions. **b)** Cartoon describing (left) the approach where binary classifiers are used in a sliding-window setting to make predictions at higher nucleotide resolution, contrasting with (right) our segmentation methods that directly make single-nucleotide predictions. **b)** Area under the precision-recall curve (auPRC) performance for sliding two binary promoter classifiers (DeePromoter and NTv2-promoters) or segmenting promoter elements with the different segmentation models. **c)** Area under the precision-recall curve (auPRC) performance for sliding the binary enhancer classifier NTv2 or segmenting enhancer elements with the different segmentation models. **d)** Inference times on a 30kb sequence for the different models. Inference times were calculated in a single A100 GPU.

SegmentNT-30kb outperformed the sliding window baselines for both promoter and enhancer annotation (Fig. 6b,c). Similar to our previous analyses (Fig. 3b), using Enformer and Borzoi as DNA encoders further improved the performance on this task, with SegmentEnformer achieving the highest performance (Fig. 6b,c). Despite the strong performance of DeePromoter on their curated test set, we observed poor generalization to the context of genomics sequences when compared with sliding the Nucleotide Transformer models, likely due to the design of their test set (see details in Methods). In addition to improved performance, since our segmentation models make all predictions at once for all nucleotides they are much faster on inference on large sequences, which will allow for faster evaluation on regions with candidate genetic variants and along full personalized genomes (Fig. 6d). These results show that our models show improved performance on annotating regulatory elements with nucleotide precision, despite the less-strict boundaries of these elements that prevent higher performance scores.

### Generalization of SegmentNT across species

We next explored how SegmentNT trained on human genomic elements could generalize to other species (Fig. 7a). Gene annotations for more distant, less-studied species are less accurate, while annotations of regulatory elements such as promoters and enhancers are very scarce. Thus, models that can predict these elements for different species hold great potential. In addition, comparison of predictions across species should provide insights about the evolutionary constraints of each element.

**Figure 7.**
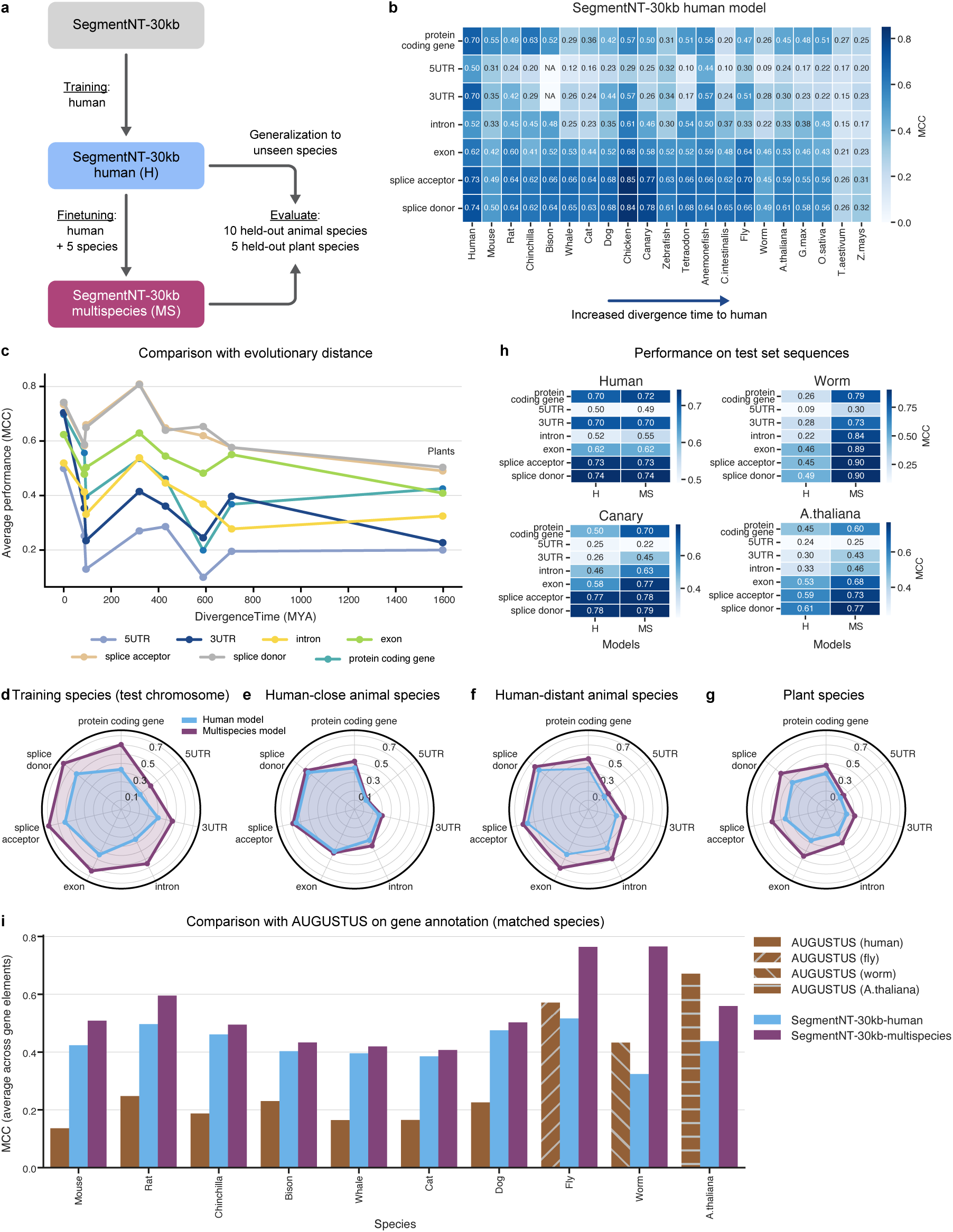
SegmentNT generalizes across species. **a)** Cartoon explaining the fine-tuning of SegmentNT models and the evaluation across unseen species. **b)** Performance of the human model on the gene elements of all species, sorted by divergence time to human. We used as metric the MCC. Data is shown as the mean MCC values from 10 different samplings of each species’ test set. **c)** Comparison between MCC performance and divergence time per gene element. The MCC was averaged for species with the same evolutionary distance. **d-g)** Radar plot depicting the performance of the human and multispecies SegmentNT models per element for **(d)** species in the training set, **(e)** human-close animal species in the test set, **(f)** human-distant animal species in the test set and **(g)** plant species in the test set. **h)** Performance of the human (H) and multispecies (MS) model per element for 4 representative species. Data is shown as the mean MCC values from 10 different samplings of each species’ test set. **i)** Performance of AUGUSTUS compared with SegmentNT-30kb human and multispecies models for the annotation of genes in the genomes of the different species. Average MCC across gene elements.

For this analysis, we selected an additional set of 15 animal and 5 plant species and for each curated a dataset of annotations for the 7 main genomic elements available from Ensembl [46], namely protein-coding gene, 5’UTR, 3’UTR, intron, exon, splice acceptor and donor sites (see Methods). This allows us to evaluate the performance of the human model in each species on the 7 element types, while for the other 7 elements our predictions might be informative of potential regulatory regions. While some of these species are less studied and thus have annotations that are mostly based on bioinformatics predictions, they still allow for the comparison of the generalization of our models against competitive approaches. Similar to the human datasets, each dataset was split in train, validation and test chromosomes, removing orthologous sequences to ensure no data leakage. We selected our best model trained on the human 14 genomic elements, SegmentNT-30kb, and evaluated it on each species test set.

We observed high performance of the human SegmentNT-30kb model across species, and particularly high for exon and splice sites, correlating with their high evolutionary conservation (Fig. 7b,c). For the other elements the performance was good for related species like gorilla and macaque, but dropped for more evolutionary-distant animals and plants. This shows that the SegmentNT-30kb model can generalize to some extent to other species, even for plants whose genome structure is very different, but that the performance depends on the evolutionary distance of the genomic elements and species.

### Multispecies SegmentNT model shows improved species generalization

Since gene elements have evolved and therefore their sequence determinants might differ between species, we trained an additional, multispecies model (SegmentNT-30kb-multispecies) by fine-tuning the human SegmentNT-30kb model on the genic annotations of human together with 5 selected animal species: mouse, chicken, fly, zebrafish and worm (see Methods). The remaining 10 animal and 5 plant species were kept as held-out test set species for comparing the generalization capabilities of the human and multispecies models. We note that since most training species have limited annotation of regulatory elements, we focused this multispecies model only on genic elements and therefore it should not be used for the prediction of regulatory elements. The performance of the SegmentNT-30kb-multispecies model improved quickly during fine-tuning, leveraging its previously acquired knowledge of human elements. We observed improved performance for the test chromosomes of the training species for the SegmentNT-30kb-multispecies model over the human SegmentNT-30kb model (Fig. 7d and Supplementary Fig. 7, 8), showing that gene elements diverged between species and it is necessary to adjust the model accordingly.

We next evaluated both human and multispecies SegmentNT-30kb models on the held-out set of 10 animal species, splitting them in two groups: 6 with an estimated divergence time from human of less than 100 million years (human-close species) and 4 more distant (more than 100 million years; human-distant; data from TimeTree). The human model generalizes well for unseen species and showed better performance for human-close (average MCC of 0.62) than human-distant species (average MCC of 0.49; Fig. 7e,f). SegmentNT-30kb-multispecies demonstrated similarly good performance on human-close species (average MCC of 0.64) and improved performance on human-distant species (average MCC of 0.57) over the human model (0.49; Fig. 7e,f). In addition, the SegmentNT-30kb-multispecies model showed improved performance over its human counterpart for the prediction of splicing variants (Supplementary Fig. 6d).

Finally, we evaluated the performance of both models on 5 plant species: *Arabidopsis thaliana*, *Glycine max* (soybean), *Oryza sativa* (rice), *Triticum aestivum* (wheat) and *Zea mays* (corn/maize). We note that the multispecies model was only trained on animal genomes and did not see any plant genome. Still, we observed a strong improvement of the SegmentNT-30kb-multispecies model over the human model across all plants (average MCC of 0.45 vs 0.34; Fig. 7g and Supplementary Fig. 8). Although the performance on plant genomes (average MCC of 0.45) was lower than on human-distant animal genomes (0.57), the SegmentNT-30kb-multispecies model can still be very useful to annotate the genomes of poorly characterised plants. This is particularly encouraging, given the large difference in genome structure between animals and plants, whose genomes are characterized by distinct evolutionary patterns including distinct genome conservation (i.e. coding versus non-coding sequence maintenance), genome architecture (e.g., repeats expansion), and notably, major polyploidization processes [47, 48]. In the case of *Zea mays* and *Triticum aestivum*, whose genomes are tetraploid and hexaploid, respectively, and have undergone large-scale rearrangements, it is encouraging that the multispecies model still retained predictive performance. Although the genomic code is not expected to change, such changes increase the sequence diversity of the different types of elements. In addition, it outperforms AUGUSTUS on gene annotation in all species except for Arabidopsis (Fig. 7i and Supplementary Fig. 9). This SegmentNT-30kb-multispecies model is thus more general and can generalize better to species not included in the training set (Fig. 7h). Altogether, these results show that SegmentNT can be easily extended to additional genomic elements and species, including plants, which opens up promising new research directions to be explored in future work.

## Discussion

Here, we introduced a novel methodology for fine-tuning DNA foundation models to segment different genic and regulatory elements at single-nucleotide resolution. Our best model, SegmentNT, is an extension of the DNA foundation model NT and is capable of processing DNA sequences up to 50kb long. We show highest performance for genic elements, including splice sites, and how each element depends on different context windows. For a given 50kb sequence, SegmentNT makes 700, 000 predictions at once allowing to annotate any input sequence in a very efficient way. SegmentNT trained on the human genome can already generalize to other species, but to make SegmentNT more broadly applicable to annotate sequences from different species we developed a multispecies version that improves generalization to unseen species. We make our best models available on our github repository and HuggingFace space.

SegmentNT provides strong evidence that DNA foundation models can tackle complex tasks in genomics at single-nucleotide resolution. Up until now, there is no consensus for the benefit of pre-trained foundation models for genomics. There has been limited improvements on most tasks where these models have been evaluated on [21–23, 26, 42, 44]. Here we focused on a more challenging task of segmenting various genomic elements in DNA sequences at nucleotide resolution, compared with single tasks of classifying short sequences as containing a given type of element. Our results show that the highest performance is achieved by combining a pre-trained NT and a segmentation U-Net head, when compared with applying such segmentation architectures directly from one-hot encoded DNA sequences. This is a strong evidence for the value added by such pre-trained models and points to the need of expanding their applications and evaluations to more realistic tasks in genomics.

A current limitation of DNA foundation models is their limited context length. NT was the pre-trained model with the largest context length at its time, trained on sequences of up to 12kb [22]. Since then different approaches have been proposed to extend the context of such models, mostly by relying on novel state-space architectures to avoid the quadratic scaling of Transformers [23, 24, 43]. Here we took a different approach and extended the context of SegmentNT through context-length extrapolation in both training and evaluation phases, showing improved performance for sequences up to 50kb (see also [42]). In addition, we showed that replacing the DNA encoder by the longer-range pre-trained genomics models Enformer [18] and Borzoi [19] allows us to segment sequences of 524kb, which improved the performance in gene regulatory elements. We expect that extending the context of NT and SegmentNT models to longer sequences with efficient context-extension approaches will yield further improvements for DNA segmentation tasks. Many techniques have recently emerged in fields like natural language processing that manage to increase the input length of Transformer models to process hundreds of thousands of tokens at a time [49–52]. These approaches together with the new developments of state-space models provide promising avenues to build the next generation of models.

We have extensively benchmarked SegmentNT against state-of-the-art tools specialized in the different domains of gene annotation, splicing detection and prediction of regulatory elements. SegmentNT generally matched or outperformed the respective competitors while being able to solve all tasks at once. For gene annotation tasks we have not compared our model with alternative approaches that consider experimental data (e.g. chromhmm [53]) or ortholog sequence alignments (e.g. CACTUS [54]) and focused on purely sequence-based tasks and baselines for a fair comparison. However, it would be worth to explore how to integrate SegmentNT within those established gene annotation pipelines.

In this work, we modeled promoter and enhancer regulatory elements based on experimentally validated genomic regions labeled by their biochemical properties, as defined by the ENCODE consortium [37]. This contrasts with approaches that directly predict biochemical marks such as chromatin accessibility, histone modifications, or transcription factor binding, as seen in models like Enformer [18] and Borzoi [19]. By considering regulatory elements as discrete annotated regions, our approach offers additional value on two main fronts: (1) such labels encapsulate multiple biochemical signals, which may collectively represent functional distinctions that are less apparent when evaluating individual marks in isolation; (2) predicting these consolidated labels allows to generalize across species better than predicting biochemical data that is more tissue/species-specific. Still, we acknowledge that the annotation we used is a simplification, given the complexity of regulatory elements in terms of their sequence features and activity across different cell types and contexts. To address this, we categorized promoters and enhancers into tissue-invariant and tissue-specific classes, already observing different performance between these groups. In future work, we expect that further refining these categories by splitting promoters and enhancers according to specific cell types, allowing the model to capture more granular, cell type-specific regulatory codes, should enhance the accuracy of regulatory element predictions.

An important result of our work is the demonstration that SegmentNT trained on human genomic elements can generalize to unseen species, both animal and plant. The generalization is stronger for splice sites and exons, likely due to their high conservation. In addition, we observed reduced generalization for species with longer divergence times to human. To improve the generalization to more distant species, we developed a SegmentNT-multispecies version that shows improved performance on unseen animal and plant species. It’s notable how this model, trained on a subset of animal species, extends its predictive ability to plant species genomes, suggesting that the sequence requirements of each genomic element captured by the model are general and can be translated to different domains. Thus, this model can be leveraged to annotate sequences up to 50kb sequences of any species *de novo* which should be useful to annotate the genomes of less-characterized species. Finally, we anticipate that further predictive power is likely to be achieved by expanding the set of species included in the multispecies model, such as incorporating plant species and particularly those with large-scale genome rearrangements to increase sequence diversity.

Overall, our work has several direct applications. First, the fine-tuned DNA encoder within SegmentNT should provide stronger representations of human genomic elements and could be used to improve performance on downstream tasks [55]. Second, interpreting the representations learned by SegmentNT could reveal insights about the genome and its encoded information. Third, the accuracy of SegmentNT predictions can be leveraged to evaluate the impact of sequence variants on the different types of genomic elements, as we showed for splicing isoforms. Thanks to the extended sequence context and the prediction of several types of genomic elements, we foresee important applications for the analysis of cancer genomes and their large structural variants. Fourth, SegmentNT-multispecies can be directly applicable to annotate and explore the genomes of different species. Fifth, SegmentNT’s architecture can be easily applied to additional genomics annotations or nucleotide-level experimental data, and combined with different DNA encoders (as demonstrated by using it with the Enformer and Borzoi models). Increasing the number of channels per nucleotides predicted by SegmentNT to include data coming from multiple experiments and biological processes should improve the transfer between tasks and lead to generalisation in a way similar to the Segment Anything Model for images [56]. We ultimately hope that SegmentNT can be a useful tool for the genomics community and foster new developments in our understanding of the genome code.

## Methods

### Genome segmentation model

In this section, we introduce our approach to segment the genome, namely SegmentNT. We formulate this problem as the segmentation of a sequence of *N* nucleotides (for example *N* = 3, 000 bp, 3kb, or *N* = 10, 000 bp, 10kb) by predicting a probability for each nucleotide to be part of one of *K* = 14 elements: *protein-coding gene*, *lncRNA*, *5’UTR*, *3’UTR*, *exon*, *intron*, *splice donor site*, *splice acceptor site*, *polyA signal*, *promoter tissue-invariant*, *promoter tissue-specific*, *enhancer tissue-invariant*, *enhancer tissue-specific* or *CTCF-bound*.

#### SegmentNT architecture

Nucleotide Transformer (NT) can be used as a backbone for segmenting a sequence of nucleotides. SegmentNT uses the pre-trained *NT-Multispecies-v2 (500M)* model as DNA encoder to extract embeddings for each of the tokens yielded by a 6-mer tokenizer. We use this model since it was the most powerful model in the NT benchmark [22]. We note *N* the number of nucleotides in the DNA sequence and *L* the number of DNA tokens (with roughly *L* ≈ *N*/6). In order to segment the sequence, we replace its original language model head by a 1-dimensional *U-Net* segmentation head [32] made of 2 downsampling convolutional blocks and 2 upsampling convolutional blocks. Each of these blocks is made of 2 convolutional layers with 2, 048 and 4, 096 kernels respectively, and *L*/2 and *L*/4 sequence length. This accounts for 63 million parameters. The purpose of adding these additional U-Net layers during fine-tuning is to better capture multi-scale, hierarchical representations along the sequence, which can be crucial for capturing fine-grained patterns in sequence data. This hierarchical feature learning improves localization and contextual awareness, as verified by improved performance when combining NTv2 with the U-Net head. In addition, this approach makes SegmentNT more flexible since it can be used with any DNA encoder independent of its output dimensions (see the examples with Enformer and Borzoi encoders). The output of the U-Net layer predicts for each nucleotide 2 logits per genomic element, with an output tensor of shape (*N*, *K*, 2). These logits are then passed through a softmax layer, applied over the last tensor dimension, that returns a tuple (*p*, 1 − *p*) per nucleotide and per genomics element where *p* is the probability of the nucleotide belonging to the element. We do not add further constraints on predictions such as the fact that one nucleotide belongs only to one element, and thus each nucleotide can be part of multiple elements. For binary classification metrics we used as threshold 0.5 to annotate nucleotides as belonging to each element type.

#### Model training and evaluation

We train our model using Adam optimizer with *lr* = 5*e* − 5. We use a batch size of 256 and trained the SegmentNT-3kb model for 10.24B tokens, meaning a total of 20.48M sequences seen during training. The training was done on a cluster of 8 GPU H100 over 20 hours. The 10kb, 20kb and 30kb models were initialized from the best checkpoint of the respective smaller model for faster adaptation to longer lengths. For example, SegmentNT-30kb model was initialized with the best SegmentNT-20kb checkpoint and fine-tuned for an additional 2.56B tokens (0.51M sequences). We use focal loss [33] with *γ* = 2 which helps the model to focus on “harder” samples, *ie* the sparse nucleotides that belong to an element.

We split our dataset between train, validation and test sets by chromosome. Namely, chromosomes 20 and 21 are used for test, chromosome 22 is used for validation, and the remaining are used for training. We excluded from the test set chunks that contain genes with homology to genes in the train or validation chromosomes, using the annotations from Ensembl BioMart (accessed 5/08/2024). During training, sequences are randomly sampled in the genome with associated annotations. We keep the sequences in the validation and test sets fixed by using a sliding window of length *N* over the respective chromosomes. The validation set was used to monitor training and for early stopping while the test set was used to evaluate model performance. We used Matthews correlation coefficient (MCC) as a validation metric and selected the best checkpoint based on the average score across all 14 genomic elements. During evaluation and testing, we predict 14 probabilities for each nucleotide in a sequence, corresponding to the different element types. We combine these predictions across all sequences into a single array for each element type and calculate performance metrics based on the predictions for every nucleotide across the dataset. These metrics treat each nucleotide as a separate prediction and include the Matthews correlation coefficient (MCC), area under the precision-recall curve (auPRC), Jaccard similarity and the F1-score.

#### Model ablations and baselines

SegmentNT is made of a DNA encoder (*NT-Multispecies-v2 (500M)*) and a 1-dimensional U-Net segmentation head, as described above. To evaluate the added value of using a pre-trained backbone encoder, we performed different model ablations and compared it on the 3kb dataset with the following models. (1) The SegmentNT where we use the Nucleotide Transformer v1 2.5B 1000G model [22] as backbone (this is the only model in this paper where the backbone is not the NT-Multispecies-v2 (500M)), featuring 2.6B parameters including the segmentation head. (2) The SegmentNT-3kb model (563M), initialized with the NT-Multispecies-v2 weights but where we only fine-tune the segmentation head. (3) Two versions of the SegmentNT model (563M) whose encoder is initialized with random weights, where we train either all parameters or only the segmentation head. (4) Two versions of the U-Net segmentation head alone, with 63M and 252M parameters respectively, which take one-hot encoded DNA sequences as input instead of the embeddings outputted by the DNA encoder. (5) The same segmentation head where we first pass the one-hot representation in a linear layer, which adds 3M parameters, bringing the model to 66M parameters, to upscale it to the embedding dimension before passing in the U-Net (1024 vocabulary).

We compared SegmentNT with the two state-of-the-art single-nucleotide model architectures BPNet [38] and SpliceAI [39] on the 3kb dataset, using directly one-hot encoded DNA sequences (of *L* = 3, 000) and varying their embedding size to test different model scaling. We trained two randomly initialized versions of the BPNet model with embedding sizes of 64 (original model) and 1024, bringing the number of parameters to respectively 120k and 29M. In addition, we trained three randomly initialized versions of the SpliceAI model with an embedding size of 32 to match the original model (700K parameters), and embedding sizes of 256 (44M) and 920 (573M) to upscale the model.

As with the SegmentNT models, we monitored the training by validating on sequences from chromosome 22 and selected the best checkpoint based on the highest average MCC score across the 14 types of elements. See Supplementary Table 1 for the number of sequences seen during training for all models. We present all model results in Supplementary Tables 2 and 3, and Supplementary Fig. 3.

#### Context-length Extension

Since the DNA encoder of SegmentNT is using rotary positional embeddings (RoPE) that have been trained on a maximum sequence length of 2, 048 tokens, its performance degrades very quickly when inferring on longer sequences. Several previous works have suggested adaptations to RoPE to better handle evaluation or fine-tuning on longer sequences, such as using Position Interpolation ([57, 58]) or “NTK-aware” scaled Rope [59]. More recently, [41] formalized different methods and augmented them to propose a final adaptation of RoPE to unseen lengths called YaRN. After testing the different approaches, YaRN did not introduce improvements to extending SegmentNT lengths compared to simply using “NTK-aware” RoPE. Since the latter is lighter to implement we decided to use it for extending the context of SegmentNT.

As described by Pend et al. [41], with the hidden layer set of hidden neurons denoted by *D*, and a sequence of vectors *x*_1_, …*x*_*L*_ ∈ *R^|^*^*D*|^, “NTK-aware” RoPE can be described by the following equation:

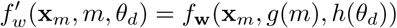

where *d* is the position along the embedding dimension, *m* is the position of the embedding in the sequence, *f* is the RoPE function (detailed in Eq.1 of [57]), *g*(*m*) = *m*, *h*(*θ*_*d*_) = *b^′−^*^2*d*/|*D*^*^|^*, 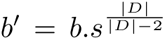 and finally 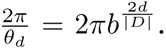 The rescaling factor *s* is computed as 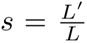 with *L^′^* the extended context length and *L* the training context length, which for the *NT-Multispecies-v2 (500M)* is 2, 048 tokens.

For SegmentNT models trained with “NTK-aware” RoPE, all sequences with length inferior to their training length are evaluated with the same rescaling factor that was used during the training. Concretely, SegmentNT-30kb is trained with *s* = 2.44, and therefore inference on a sequence smaller than 30, 000bp is done with *s* = 2.44. When evaluated on a 50kb sequence, the rescaling factor becomes *s* = 4.07.

#### Enformer and Borzoi as segmentation encoders

We trained additional segmentation models using Enformer [18] and Borzoi [19] as DNA encoders. These models contain a tower of convolution- and subsampling blocks followed by a series of self-attention blocks operating at 128bp resolution embedding vectors. Borzoi also makes use of a U-net architecture to increase the resolution back to 32bp. Both models were pre-trained in a supervised manner to predict thousands of epigenetic and gene expression tracks in various mouse and human cell types. We have re-implemented Enformer and Borzoi in our Jax codebase and have validated their exact reproduction by showing an absolute difference of less than 1e-5 in all activations as well as in the predictions between the original implementation and ours. To use Enformer and Borzoi as DNA encoders, we place the U-Net segmentation head on top of their last layer representations, before the prediction heads, and call these models SegmentEnformer and SegmentBorzoi, respectively. We fine-tuned the whole network on our segmentation dataset using either 30kb input sequences (for a fair comparison to SegmentNT-30kb) or the model’s original input length, 196kb for Enformer and 524kb for Borzoi. All training and validation hyperparameters remained the same.

#### Multi-species training

We trained an additional, multispecies model (SegmentNT-30kb-multispecies) by fine-tuning the human SegmentNT-30kb model on the annotations of human and five additional species together: mouse (*mm*10), chicken (*galGal*6), fly (*dm*6), zebrafish (*danRer*11) and worm (*ce*11). We used the same model hyperparameters and training parameters. Since the different species have different genome sizes, we balanced examples from each dataset with the following weights: 5 for human, 4 for mouse, 2 for chicken, fly and zebrafish, and 1 for worm. Similar to the human dataset, we held-out specific chromosomes for validation and testing: mouse (validation: chr19, test: chr18), chicken (validation: chr28, test: chr16 and chr27), fly (validation: chr4, test: chr2L), zebrafish (validation: chr25, test: chr23 and chr24), worm (validation: chrI, test: chrIII).

### Genome annotation data

#### Human genomic elements

The human segmentation dataset of genomic elements was created from 14 types of elements, divided in gene elements (protein-coding genes, lncRNAs, 5’UTR, 3’UTR, exon, intron, splice acceptor and donor sites) and regulatory elements (polyA signal, tissue-invariant and tissue-specific promoters and enhancers, and CTCF-bound sites). The final segmentation dataset was created by overlapping all 14 elements with every DNA sequence of length *N* nucleotides. Sequences with Ns were removed.

The location of all gene elements and polyA signals were obtained from GENCODE [35] V44 gene annotation. Annotations were filtered to exclude level 3 transcripts (automated annotation), so all training data was annotated by a human. We used *extract_splice_sites.py* from HISAT2 [60] (https://github.com/DaehwanKimLab/hisat2/blob/master/hisat2_extract_splice_sites.py) to extract respective intron and splice site annotations.

Promoter, enhancer and CTCF-bound sites were retrieved from ENCODE’s SCREEN database (https://screen.wenglab.org/) [37]. Distal and proximal enhancers were combined. Promoters and enhancers were split in tissue-invariant and tissue-specific based on the vocabulary from Wouter Meuleman et al. [61] https://www.meuleman.org/research/dhsindex/. Enhancers or promoters overlapping regions classified as tissue-invariant were defined as that, while all other enhancers and promoters were defined as tissue-specific.

#### Multi-species dataset

To create segmentation datasets for additional species we focused only on the main gene elements: protein-coding genes, 5’UTR, 3’UTR, exon, intron, splice acceptor and donor sites. We obtained their annotations as described for the human dataset but retrieved from Ensembl databases (https://www.ensembl.org). We considered 5 species to train the multispecies model: mouse (*mm*10), chicken (*galGal*6), fly (*dm*6), zebrafish (*danRer*11) and worm (*ce*11). We created a held-out test set made of 10 animal species: bison (*Bison*_*U MD*1), whale (*ASM* 228892*v*3), cat (*F elis*_*catus*_9), dog (*ROS*_*Cfam*_1), canary (*SCA*1), tetraodon (*T ET RAODON* 8), anemonefish (*AmpOce*1), rat (*mRatBN* 7), chinchilla (*ChiLan*1) and Ciona intestinalis (*KH*). We added a second held-out test set of 5 plant species: Arabidopsis thaliana (*T AIR*10), Glycine max (soybean, *Glycine*_*max*_*v*2.1), Oryza sativa (rice, (*IRGSP* 1.0), Triticum aestivum (wheat, *IW GSC*) and Zea mays (corn/maize, *Zm*_*B*73_*REF ERENCE*_*NAM*_5.0). We excluded from each species’ test set any chunk that contained genes that show orthology to genes in the human train or validation chromosomes, using the annotations from Ensembl BioMart (accessed 5/08/2024). Evolutionary distance data was retrieved from Timetree of Life.

### Benchmarking for regulatory elements

#### Sliding Nucleotide Transformer fine-tuned models

We compared our segmentation models (SegmentNT-30kb, SegmentEnformer-30kb and SegmentBorzoi-30kb) with a sliding window approach, where a binary classifier is used to predict the output probability for multiple short sliding windows over the whole test chromosomes. We applied this approach for the segmentation of promoters and enhancers, combining tissue-invariant and tissue-specific classes into a single promoter and enhancer class, since common baselines were trained to predict promoters and enhancers globally. As binary classifier we used the NT-v2 fine-tuned models on these promoter and enhancer regions, respectively, as they performed the best in the NT benchmark [22]. Sliding windows were created using a step size of 10 and the input size of the respective promoter (300bp) and enhancer (400bp) models. The 10bp step size was used for a compromise between high resolution but limited number of windows per genome region. The performance was evaluated using auPRC. All inference times were calculated in a single A100 GPU.

#### Sliding DeePromoter

We evaluated the performance of DeePromoter [7] for predicting promoter elements along the genome. Since we could not access the web-server for promoter prediction developed by the authors (https://home.jbnu.ac.kr/NSCL/deepromoter.htm; accessed 09/09/2024), we retrained their model using the pytorch implementation provided at https://github.com/egochao/DeePromoter/. We reproduced the results in their test set with high performance (MCC for human TATA promoters: 0.82, non-TATA promoters: 0.89; compared with 0.88 and 0.92 reported in the original paper [7]), ensuring that we have accurate model checkpoints to evaluate on our benchmark. Similar to the NT baseline, we evaluated DeePromoter by sliding the model over the whole test chromosomes and evaluated if promoter regions are correctly predicted by the model over background regions. We combined both TATA and non-TATA DeePromoter models and used as prediction values the highest of the two per window. Sliding windows were created using a step size of 10 and the input size of the respective models (300bp). The performance was evaluated using auPRC. All inference times were calculated in a single A100 GPU.

### Benchmark on gene annotation

We compared the gene annotation capabilities of SegmentNT with the state-of-the-art gene finder AUGUSTUS [5, 6] in three different settings, including segments of only genes and predicting only the main isoform or all isoforms, or in whole-chromosome setting. We first evaluated the models in the gene annotation task presented at the recent BEND benchmark [44], analysing only sequences in the SegmentNT test chromosomes 20 and 21. We have adapted it to create a version with windows of 30kb that contain genes with only a single isoform, with a single label per nucleotide, and a second version that contains all genes and respective isoforms, allowing multiple labels per nucleotide. The third dataset was the actual test set of SegmentNT, where the model needs to annotate the gene regions within the whole test set chromosomes and including all their annotated isoforms. We created this dataset for human but also all other species where AUGUSTUS could be tested: these include all mammals, where we used the human AUGUSTUS as recommended by the authors; and other species with specialized models (fly, worm and Arabidopsis). We measured the annotation performance on all datasets using the standard F1-score and the MCC metric, using 0.5 as the probability threshold, and analysed their precision and recall values.

Augustus was run with the following settings for the single isoform setting (example for human genome):

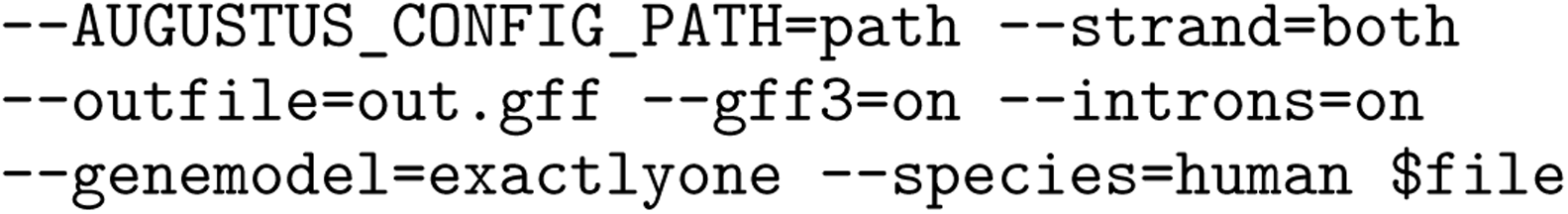

and with the following settings for the multiple isoform setting (example for human genome):

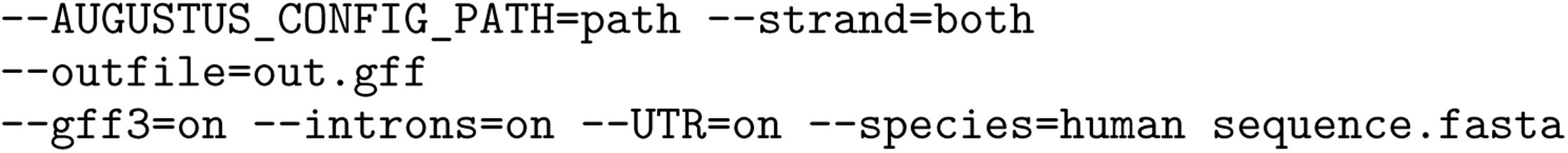

### Benchmarking on splicing tasks

#### Comparison with SpliceAI and Pangolin

We compared SegmentNT with SpliceAI [39] and Pangolin [45] on both SpliceAI’s test set and SegmentNT’s test set given their different settings. We used the scripts available at the Illumina Basespace platform ^1^ to reproduce the testing dataset presented in SpliceAI and adapted it for a fair comparison with SegmentNT. Namely, we consider sequence windows of 30kb (instead of 10kb in the original publication) and compute the predictions on the whole window (instead of not predicting for the flanking +/- 5kb sequence). We have also removed sequences with Ns due to the constraint of the SegmentNT architecture. This test set contains only mRNA sequences and all in the forward strand (i.e. for genes in the reverse strand, the sequence is reversed to have the gene in the forward orientation). We also compared both models on the SegmentNT’s 30kb human and multispecies test sets, where we filtered for windows that only contain genes in the forward strand, given the training settings of SpliceAI and Pangolin. Since Pangolin does not differentiate between acceptor and donor sites, predicting a single splice site label, we have converted these predictions into acceptor and donor predictions for a more direct comparison. More specifically, when calculating the splice acceptor predictions, we compare Pangolin splice site predictions on acceptor sites versus the remaining nucleotides after removing annotated donor sites; for donor sites we do the opposite, comparing Pangolin predictions on donor sites versus the remaining nucleotides after removing annotated acceptor sites. We used as performance metrics both auPRC, MCC and top-k [39]. For the comparison across multiple species, we averaged the performance of each model in each genomic element type across species.

## Data availability

The SegmentNT training data was obtained from publicly available resources. Gene annotations were obtained from GENCODE (https://www.gencodegenes.org/) and Ensembl databases (https://www. ensembl.org). Human regulatory elements were obtained from ENCODE’s SCREEN database (https://screen.wenglab.org/). Evolutionary distance data was retrieved from Timetree of Life. SpliceAI test set data was derived from the Illumina Basespace platform (https://basespace.illumina.com/projects/ 66029966/). We have also created an interactive browser session with the labels and predictions of the human SegmentNT-30kb model along the test chromosomes 20 and 21 at https://tinyurl.com/23837bnl.

## Code availability

Model weights of the human and multispecies SegmentNT-30kb models, SegmentEnformer and SegmentBor-zoi models, as well as inference code in Jax are available for research purposes at https://github.com/instadeepai/nucleotide-transformer?tab=readme-ov-file#the-segmentnt-models. Hugging-Face versions of the models, in PyTorch, can be found at https://huggingface.co/collections/InstaDeepAI/segmentnt-65eb4941c57808b4a3fe1319. Example notebooks are available on Google Colab at https://colab.research.google.com/github/instadeepai/nucleotide-transformer/blob/main/examples/inference_segment_nt.ipynb.

### Acknowledgments

We thank Aliou Kayantao for his help improving the figure panels. We would also like to thank Volodymyr Kuleshov, Yang Li and the Kuleshov lab for insightful discussions about context-length extension and important applications for DNA foundation models. Finally, we would like to thank Lida Rosseló for help on the project management side of this research project.

## Competing interests

B.P.d.A., H.D-T., G.R., M.G., J.M-R., Z.T., D.M.E., M.L., A.L., K.B. and T.P. are employees of InstaDeep LTD. C.B., L.H., P.P., M.L. and U.S. are employees of BioNTech LTD.

## Supplementary Tables

**Supplementary Table 1.** Baselines models training details.

| Model | Pre-trained DNA encoder | Number of parameters | Number of training sequences seen (M) |
| --- | --- | --- | --- |
| BPNet arch. | No | 120k | 53 |
| BPNet arch. Large | No | 29M | 49 |
| SpliceAI arch. | No | 700k | 125 |
| SpliceAI arch. large | No | 44M | 61 |
| SpliceAI arch. extra-large | No | 573M | 45 |
| UNet one-hot | No | 63M | 14 |
| UNet 1024-vocab | No | 66M | 20 |
| UNet large one-hot | No | 252M | 18 |
| SegmentNT-3kb (NTv1 human; 2.5B) | NTv1-1000G (2.5B) | 2.6B | 22 |
| SegmentNT-3kb (random-init) | No | 563M | 79 |
| SegmentNT-3kb (random-init; only head) | No | 563M | 18 |
| SegmentNT-3kb (only head) | NTv2 500M | 563M | 12 |
| SegmentNT-3kb | NTv2 500M | 563M | 20 |
| SegmentEnformer | Enformer | 405M | 14 |
| SegmentBorzoi | Borzoi | 340M | 14 |

**Supplementary Table 2.** Performance of all models across the 14 genomic elements (MCC). We used as metric the Matthews correlation coefficient (MCC). Data are presented as mean MCC values +/- standard deviation from 10 different samplings of the test set.

| Model | 3UTR | 5UTR | CTCF-bound | enhancer tissue-invariant | enhancer tissue-specific | exon | intron |
| --- | --- | --- | --- | --- | --- | --- | --- |
| BPNet arch. | 0.00 (± 0.002) | 0.17 (± 0.003) | 0.00 (± 0.000) | 0.00 (± 0.000) | 0.04 (± 0.001) | 0.24 (± 0.002) | 0.22 (± 0.001) |
| BPNet arch. large | 0.10 (± 0.004) | 0.26 (± 0.004) | 0.06 (± 0.003) | 0.12 (± 0.003) | 0.20 (± 0.001) | 0.27 (± 0.001) | 0.21 (± 0.001) |
| SpliceAI arch. | 0.27 (± 0.009) | 0.23 (± 0.006) | -0.0 (± 0.000) | 0.00 (± 0.002) | 0.13 (± 0.002) | 0.38 (± 0.003) | 0.31 (± 0.002) |
| SpliceAI arch. large | 0.27 (± 0.006) | 0.30 (± 0.006) | 0.06 (± 0.004) | 0.17 (± 0.008) | 0.26 (± 0.001) | 0.37 (± 0.005) | 0.28 (± 0.003) |
| SpliceAI arch. extra-large | 0.25 (± 0.006) | 0.29 (± 0.003) | 0.01 (± 0.005) | 0.21 (± 0.003) | 0.20 (± 0.001) | 0.37 (± 0.002) | 0.20 (± 0.003) |
| UNet | -0.0 (± 0.001) | 0.08 (± 0.001) | 0.00 (± 0.000) | 0.03 (± 0.001) | 0.02 (± 0.000) | 0.17 (± 0.001) | 0.18 (± 0.001) |
| UNet 1024-vocab | 0.00 (± 0.000) | 0.09 (± 0.001) | 0.00 (± 0.001) | 0.04 (± 0.001) | 0.03 (± 0.000) | 0.14 (± 0.001) | 0.17 (± 0.001) |
| UNet large | 0.01 (± 0.000) | 0.13 (± 0.002) | 0.02 (± 0.000) | 0.05 (± 0.001) | 0.09 (± 0.000) | 0.16 (± 0.000) | 0.12 (± 0.001) |
| Random-Init (only head) | -0.0 (± 0.000) | 0.00 (± 0.000) | 0.00 (± 0.000) | -0.0 (± 0.000) | 0.01 (± 0.000) | 0.03 (± 0.001) | 0.06 (± 0.001) |
| Random-Init | 0.14 (± 0.007) | 0.23 (± 0.007) | 0.00 (± 0.002) | 0.05 (± 0.005) | 0.14 (± 0.002) | 0.27 (± 0.004) | 0.20 (± 0.004) |
| SegmentNT-3kb (only head) | 0.34 (± 0.004) | 0.41 (± 0.008) | 0.02 (± 0.001) | 0.16 (± 0.005) | 0.18 (± 0.002) | 0.48 (± 0.002) | 0.28 (± 0.001) |
| SegmentNT-3kb (NTv1 human; 2.5B) | 0.42 (± 0.015) | 0.38 (± 0.006) | 0.08 (± 0.006) | 0.13 (± 0.006) | 0.20 (± 0.002) | 0.48 (± 0.006) | 0.29 (± 0.003) |
| SegmentNT-3kb | 0.51 (± 0.015) | 0.44 (± 0.008) | 0.08 (± 0.004) | 0.19 (± 0.004) | 0.25 (± 0.001) | 0.52 (± 0.005) | 0.30 (± 0.003) |
| SegmentNT-3kb-single-task (x14) | 0.49 (± 0.013) | 0.42 (± 0.006) | 0.09 (± 0.002) | 0.17 (± 0.006) | 0.29 (± 0.001) | 0.55 (± 0.003) | 0.33 (± 0.003) |
| SegmentNT-10kb | 0.66 (± 0.007) | 0.49 (± 0.007) | 0.09 (± 0.003) | 0.19 (± 0.007) | 0.27 (± 0.001) | 0.59 (± 0.003) | 0.40 (± 0.004) |
| SegmentNT-20kb | 0.69 (± 0.009) | 0.49 (± 0.007) | 0.09 (± 0.005) | 0.18 (± 0.009) | 0.27 (± 0.002) | 0.62 (± 0.003) | 0.47 (± 0.008) |
| SegmentNT-30kb | 0.70 (± 0.007) | 0.50 (± 0.003) | 0.08 (± 0.002) | 0.19 (± 0.005) | 0.27 (± 0.002) | 0.62 (± 0.003) | 0.52 (± 0.005) |
| SegmentEnformer-30kb | 0.65 (± 0.008) | 0.40 (± 0.006) | 0.17 (± 0.006) | 0.29 (± 0.007) | 0.44 (± 0.002) | 0.59 (± 0.004) | 0.49 (± 0.011) |
| SegmentEnformer-196kb | 0.68 (± 0.039) | 0.44 (± 0.015) | 0.19 (± 0.014) | 0.28 (± 0.023) | 0.42 (± 0.005) | 0.56 (± 0.015) | 0.60 (± 0.017) |
| SegmentBorzo-30kb | 0.68 (± 0.011) | 0.43 (± 0.008) | 0.12 (± 0.007) | 0.25 (± 0.008) | 0.42 (± 0.001) | 0.59 (± 0.004) | 0.46 (± 0.007) |
| SegmentBorzo-524kb | 0.85 (± 0.127) | 0.60 (± 0.068) | 0.16 (± 0.023) | 0.24 (± 0.050) | 0.37 (± 0.022) | 0.49 (± 0.089) | 0.52 (± 0.059) |

**Supplementary Table 2** | Performance of all models across the 14 genomic elements (MCC). We used as metric the Matthews correlation coefficient (MCC). Data are presented as mean MCC values ± standard deviation from 10 different samplings of the test set.
| Model | lncRNA | polyA signal | promoter tissue-invariant | promoter tissue-specific | protein coding gene | splice acceptor | splice donor |
| --- | --- | --- | --- | --- | --- | --- | --- |
| BPNet arch. | 0.00 (± 0.000) | 0.00 (± 0.000) | 0.26 (± 0.005) | 0.05 (± 0.003) | 0.30 (± 0.001) | 0.03 (± 0.004) | 0.05 (± 0.002) |
| BPNet arch. large | 0.00 (± 0.000) | 0.04 (± 0.005) | 0.40 (± 0.005) | 0.06 (± 0.004) | 0.29 (± 0.001) | 0.22 (± 0.003) | 0.29 (± 0.002) |
| SpliceAI arch. | 0.00 (± 0.000) | 0.02 (± 0.006) | 0.41 (± 0.007) | 0.05 (± 0.008) | 0.42 (± 0.003) | 0.09 (± 0.005) | 0.22 (± 0.003) |
| SpliceAI arch. large | 0.00 (± 0.002) | 0.16 (± 0.006) | 0.50 (± 0.008) | 0.13 (± 0.013) | 0.40 (± 0.002) | 0.38 (± 0.003) | 0.50 (± 0.003) |
| SpliceAI arch. extra-large | 0.01 (± 0.002) | 0.12 (± 0.006) | 0.53 (± 0.007) | 0.21 (± 0.007) | 0.20 (± 0.001) | 0.45 (± 0.002) | 0.52 (± 0.003) |
| UNet | 0.00 (± 0.000) | 0.00 (± 0.004) | 0.13 (± 0.002) | 0.01 (± 0.001) | 0.14 (± 0.001) | 0.08 (± 0.002) | 0.19 (± 0.003) |
| UNet 1024-vocab | -0.0 (± 0.000) | 0.00 (± 0.002) | 0.14 (± 0.003) | 0.00 (± 0.001) | 0.13 (± 0.001) | 0.08 (± 0.002) | 0.20 (± 0.002) |
| UNet large | -0.0 (± 0.001) | 0.02 (± 0.002) | 0.23 (± 0.004) | 0.01 (± 0.000) | 0.12 (± 0.001) | 0.22 (± 0.002) | 0.28 (± 0.002) |
| Random-Init (only head) | -0.0 (± 0.000) | -0.0 (± 0.001) | 0.00 (± 0.001) | 0.00 (± 0.000) | 0.06 (± 0.000) | 0.00 (± 0.001) | -0.0 (± 0.002) |
| Random-Init | 0.04 (± 0.006) | 0.09 (± 0.006) | 0.40 (± 0.011) | 0.13 (± 0.013) | 0.26 (± 0.004) | 0.09 (± 0.003) | 0.19 (± 0.004) |
| SegmentNT-3kb (only head) | 0.01 (± 0.001) | 0.25 (± 0.011) | 0.54 (± 0.006) | 0.20 (± 0.006) | 0.40 (± 0.001) | 0.67 (± 0.003) | 0.70 (± 0.002) |
| SegmentNT-3kb (NTv1 human; 2.5B) | 0.04 (± 0.004) | 0.28 (± 0.006) | 0.56 (± 0.007) | 0.22 (± 0.010) | 0.41 (± 0.002) | 0.62 (± 0.002) | 0.64 (± 0.003) |
| SegmentNT-3kb | 0.05 (± 0.004) | 0.30 (± 0.006) | 0.58 (± 0.006) | 0.22 (± 0.011) | 0.42 (± 0.003) | 0.69 (± 0.004) | 0.70 (± 0.002) |
| SegmentNT-3kb-single-task (x14) | 0.02 (± 0.004) | 0.34 (± 0.005) | 0.51 (± 0.005) | 0.16 (± 0.006) | 0.43 (± 0.004) | 0.71 (± 0.003) | 0.71 (± 0.003) |
| SegmentNT-10kb | 0.06 (± 0.013) | 0.36 (± 0.007) | 0.58 (± 0.009) | 0.23 (± 0.009) | 0.58 (± 0.008) | 0.71 (± 0.006) | 0.73 (± 0.005) |
| SegmentNT-20kb | 0.07 (± 0.012) | 0.38 (± 0.008) | 0.59 (± 0.009) | 0.25 (± 0.007) | 0.66 (± 0.006) | 0.71 (± 0.003) | 0.73 (± 0.003) |
| SegmentNT-30kb | 0.06 (± 0.014) | 0.40 (± 0.004) | 0.58 (± 0.007) | 0.23 (± 0.006) | 0.70 (± 0.003) | 0.73 (± 0.002) | 0.74 (± 0.003) |
| SegmentEnformer-30kb | 0.13 (± 0.011) | 0.00 (± 0.000) | 0.65 (± 0.011) | 0.33 (± 0.008) | 0.68 (± 0.009) | 0.00 (± 0.000) | 0.00 (± 0.000) |
| SegmentEnformer-196kb | 0.20 (± 0.025) | 0.01 (± 0.015) | 0.59 (± 0.042) | 0.26 (± 0.025) | 0.83 (± 0.018) | 0.00 (± 0.000) | 0.00 (± 0.000) |
| SegmentBorzo-30kb | 0.17 (± 0.011) | 0.18 (± 0.007) | 0.64 (± 0.008) | 0.29 (± 0.006) | 0.65 (± 0.007) | 0.00 (± 0.012) | 0.00 (± 0.006) |
| SegmentBorzo-524kb | 0.19 (± 0.121) | 0.16 (± 0.031) | 0.28 (± 0.306) | 0.33 (± 0.064) | 0.88 (± 0.064) | 0.04 (± 0.046) | 0.08 (± 0.035) |

**Supplementary Table 3.** Performance of all models across the 14 genomic elements (auPRC). We used as metric the area under the precision recall curve (auPRC). Data are presented as mean auPRC values +/- standard deviation from 10 different samplings of the test set.

| Model | 3UTR | 5UTR | CTCF-bound | enhancer tissue-invariant | enhancer tissue-specific | exon | intron |
| --- | --- | --- | --- | --- | --- | --- | --- |
| BPNet arch. | 0.06 (± 0.002) | 0.15 (± 0.002) | 0.01 (± 0.000) | 0.04 (± 0.001) | 0.23 (± 0.001) | 0.24 (± 0.002) | 0.62 (± 0.001) |
| BPNet arch. large | 0.07 (± 0.002) | 0.21 (± 0.003) | 0.06 (± 0.001) | 0.11 (± 0.002) | 0.30 (± 0.000) | 0.26 (± 0.001) | 0.62 (± 0.001) |
| SpliceAI arch. | 0.23 (± 0.006) | 0.21 (± 0.004) | 0.02 (± 0.000) | 0.05 (± 0.001) | 0.29 (± 0.001) | 0.37 (± 0.003) | 0.69 (± 0.001) |
| SpliceAI arch. large | 0.22 (± 0.007) | 0.21 (± 0.006) | 0.07 (± 0.002) | 0.12 (± 0.003) | 0.33 (± 0.001) | 0.35 (± 0.005) | 0.67 (± 0.001) |
| SpliceAI arch. extra-large | 0.18 (± 0.006) | 0.21 (± 0.003) | 0.07 (± 0.001) | 0.14 (± 0.002) | 0.31 (± 0.001) | 0.34 (± 0.002) | 0.59 (± 0.001) |
| UNet | 0.02 (± 0.000) | 0.08 (± 0.001) | 0.02 (± 0.000) | 0.03 (± 0.000) | 0.17 (± 0.001) | 0.15 (± 0.001) | 0.52 (± 0.001) |
| UNet 1024-vocab | 0.01 (± 0.000) | 0.05 (± 0.001) | 0.01 (± 0.000) | 0.03 (± 0.000) | 0.17 (± 0.000) | 0.12 (± 0.001) | 0.51 (± 0.001) |
| UNet large | 0.01 (± 0.000) | 0.06 (± 0.001) | 0.01 (± 0.000) | 0.02 (± 0.000) | 0.17 (± 0.000) | 0.13 (± 0.001) | 0.51 (± 0.001) |
| Random-Init (only head) | 0.01 (± 0.000) | 0.00 (± 0.000) | 0.01 (± 0.000) | 0.00 (± 0.000) | 0.08 (± 0.000) | 0.05 (± 0.000) | 0.48 (± 0.000) |
| Random-Init | 0.08 (± 0.003) | 0.13 (± 0.004) | 0.01 (± 0.000) | 0.02 (± 0.001) | 0.18 (± 0.001) | 0.23 (± 0.004) | 0.59 (± 0.002) |
| SegmentNT-3kb (only head) | 0.29 (± 0.003) | 0.33 (± 0.006) | 0.03 (± 0.000) | 0.10 (± 0.002) | 0.27 (± 0.001) | 0.44 (± 0.001) | 0.68 (± 0.001) |
| SegmentNT-3kb (NTv1 human; 2.5B) | 0.36 (± 0.014) | 0.31 (± 0.005) | 0.04 (± 0.002) | 0.10 (± 0.003) | 0.30 (± 0.002) | 0.45 (± 0.005) | 0.69 (± 0.001) |
| SegmentNT-3kb | 0.43 (± 0.010) | 0.38 (± 0.007) | 0.03 (± 0.001) | 0.10 (± 0.003) | 0.30 (± 0.001) | 0.48 (± 0.005) | 0.71 (± 0.001) |
| SegmentNT-3kb-single-task (x14) | 0.46 (± 0.005) | 0.36 (± 0.005) | 0.03 (± 0.001) | 0.10 (± 0.003) | 0.32 (± 0.001) | 0.53 (± 0.002) | 0.73 (± 0.001) |
| SegmentNT-10kb | 0.64 (± 0.008) | 0.43 (± 0.009) | 0.04 (± 0.001) | 0.10 (± 0.004) | 0.31 (± 0.001) | 0.56 (± 0.003) | 0.79 (± 0.003) |
| SegmentNT-20kb | 0.70 (± 0.008) | 0.44 (± 0.007) | 0.04 (± 0.001) | 0.09 (± 0.003) | 0.30 (± 0.001) | 0.58 (± 0.003) | 0.81 (± 0.003) |
| SegmentNT-30kb | 0.70 (± 0.006) | 0.44 (± 0.004) | 0.04 (± 0.001) | 0.10 (± 0.004) | 0.29 (± 0.002) | 0.58 (± 0.004) | 0.84 (± 0.002) |
| SegmentEnformer-30kb | 0.67 (± 0.011) | 0.37 (± 0.006) | 0.16 (± 0.003) | 0.27 (± 0.005) | 0.53 (± 0.003) | 0.57 (± 0.006) | 0.83 (± 0.004) |
| SegmentEnformer-196kb | 0.65 (± 0.051) | 0.37 (± 0.020) | 0.09 (± 0.008) | 0.16 (± 0.020) | 0.45 (± 0.006) | 0.50 (± 0.020) | 0.88 (± 0.010) |
| SegmentBorzo-30kb | 0.67 (± 0.010) | 0.39 (± 0.010) | 0.15 (± 0.003) | 0.22 (± 0.005) | 0.50 (± 0.003) | 0.60 (± 0.005) | 0.83 (± 0.002) |
| SegmentBorzo-524kb | 0.90 (± 0.103) | 0.55 (± 0.093) | 0.08 (± 0.010) | 0.13 (± 0.038) | 0.38 (± 0.025) | 0.41 (± 0.096) | 0.81 (± 0.055) |

**Supplementary Table 3** | Performance of all models across the 14 genomic elements (auPRC). We used as metric the area under the precision recall curve (auPRC). Data are presented as mean auPRC values ± standard deviation from 10 different samplings of the test set.
| Model | lncRNA | polyA signal | promoter tissue-invariant | promoter tissue-specific | protein coding gene | splice acceptor | splice donor |
| --- | --- | --- | --- | --- | --- | --- | --- |
| BPNet arch. | 0.22 (± 0.001) | 0.00 (± 0.000) | 0.29 (± 0.006) | 0.10 (± 0.002) | 0.59 (± 0.001) | 0.02 (± 0.000) | 0.02 (± 0.000) |
| BPNet arch. large | 0.21 (± 0.001) | 0.03 (± 0.001) | 0.45 (± 0.007) | 0.12 (± 0.004) | 0.57 (± 0.001) | 0.21 (± 0.001) | 0.28 (± 0.001) |
| SpliceAI arch. | 0.23 (± 0.001) | 0.01 (± 0.000) | 0.43 (± 0.009) | 0.11 (± 0.002) | 0.69 (± 0.001) | 0.06 (± 0.002) | 0.16 (± 0.004) |
| SpliceAI arch. large | 0.18 (± 0.001) | 0.09 (± 0.003) | 0.49 (± 0.010) | 0.11 (± 0.007) | 0.67 (± 0.001) | 0.35 (± 0.004) | 0.47 (± 0.003) |
| SpliceAI arch. extra-large | 0.20 (± 0.001) | 0.06 (± 0.003) | 0.49 (± 0.012) | 0.15 (± 0.004) | 0.54 (± 0.002) | 0.41 (± 0.001) | 0.49 (± 0.002) |
| UNet | 0.20 (± 0.001) | 0.01 (± 0.000) | 0.12 (± 0.003) | 0.06 (± 0.001) | 0.44 (± 0.002) | 0.07 (± 0.001) | 0.15 (± 0.003) |
| UNet 1024-vocab | 0.20 (± 0.001) | 0.00 (± 0.000) | 0.12 (± 0.004) | 0.04 (± 0.001) | 0.43 (± 0.001) | 0.06 (± 0.001) | 0.15 (± 0.002) |
| UNet large | 0.20 (± 0.001) | 0.01 (± 0.000) | 0.16 (± 0.004) | 0.04 (± 0.001) | 0.42 (± 0.001) | 0.14 (± 0.002) | 0.21 (± 0.002) |
| Random-Init (only head) | 0.17 (± 0.000) | 0.00 (± 0.000) | 0.00 (± 0.000) | 0.00 (± 0.000) | 0.37 (± 0.000) | 0.00 (± 0.000) | 0.00 (± 0.000) |
| Random-Init | 0.21 (± 0.002) | 0.03 (± 0.002) | 0.35 (± 0.015) | 0.07 (± 0.008) | 0.55 (± 0.003) | 0.06 (± 0.002) | 0.10 (± 0.003) |
| SegmentNT-3kb (only head) | 0.22 (± 0.001) | 0.16 (± 0.007) | 0.55 (± 0.006) | 0.13 (± 0.005) | 0.67 (± 0.001) | 0.58 (± 0.004) | 0.63 (± 0.003) |
| SegmentNT-3kb (NTv1 human; 2.5B) | 0.21 (± 0.002) | 0.19 (± 0.009) | 0.55 (± 0.007) | 0.17 (± 0.006) | 0.69 (± 0.001) | 0.57 (± 0.002) | 0.60 (± 0.003) |
| SegmentNT-3kb | 0.21 (± 0.001) | 0.19 (± 0.005) | 0.55 (± 0.014) | 0.14 (± 0.007) | 0.72 (± 0.002) | 0.62 (± 0.003) | 0.65 (± 0.003) |
| SegmentNT-3kb-single-task (x14) | 0.22 (± 0.001) | 0.26 (± 0.004) | 0.50 (± 0.008) | 0.11 (± 0.004) | 0.72 (± 0.001) | 0.68 (± 0.002) | 0.69 (± 0.002) |
| SegmentNT-10kb | 0.23 (± 0.007) | 0.28 (± 0.006) | 0.57 (± 0.012) | 0.15 (± 0.007) | 0.83 (± 0.004) | 0.65 (± 0.006) | 0.67 (± 0.005) |
| SegmentNT-20kb | 0.23 (± 0.005) | 0.30 (± 0.006) | 0.58 (± 0.013) | 0.16 (± 0.006) | 0.86 (± 0.002) | 0.64 (± 0.003) | 0.66 (± 0.003) |
| SegmentNT-30kb | 0.25 (± 0.003) | 0.31 (± 0.005) | 0.55 (± 0.012) | 0.15 (± 0.004) | 0.89 (± 0.001) | 0.65 (± 0.002) | 0.66 (± 0.002) |
| SegmentEnformer-30kb | 0.27 (± 0.008) | 0.01 (± 0.001) | 0.66 (± 0.012) | 0.31 (± 0.011) | 0.86 (± 0.005) | 0.01 (± 0.000) | 0.01 (± 0.000) |
| SegmentEnformer-196kb | 0.35 (± 0.017) | 0.02 (± 0.003) | 0.55 (± 0.041) | 0.17 (± 0.016) | 0.93 (± 0.009) | 0.02 (± 0.001) | 0.02 (± 0.001) |
| SegmentBorzo-30kb | 0.29 (± 0.008) | 0.15 (± 0.006) | 0.66 (± 0.007) | 0.26 (± 0.006) | 0.86 (± 0.003) | 0.10 (± 0.002) | 0.12 (± 0.003) |
| SegmentBorzo-524kb | 0.40 (± 0.100) | 0.10 (± 0.038) | 0.46 (± 0.305) | 0.22 (± 0.075) | 0.98 (± 0.021) | 0.08 (± 0.032) | 0.10 (± 0.024) |

**Supplementary Table 4.**
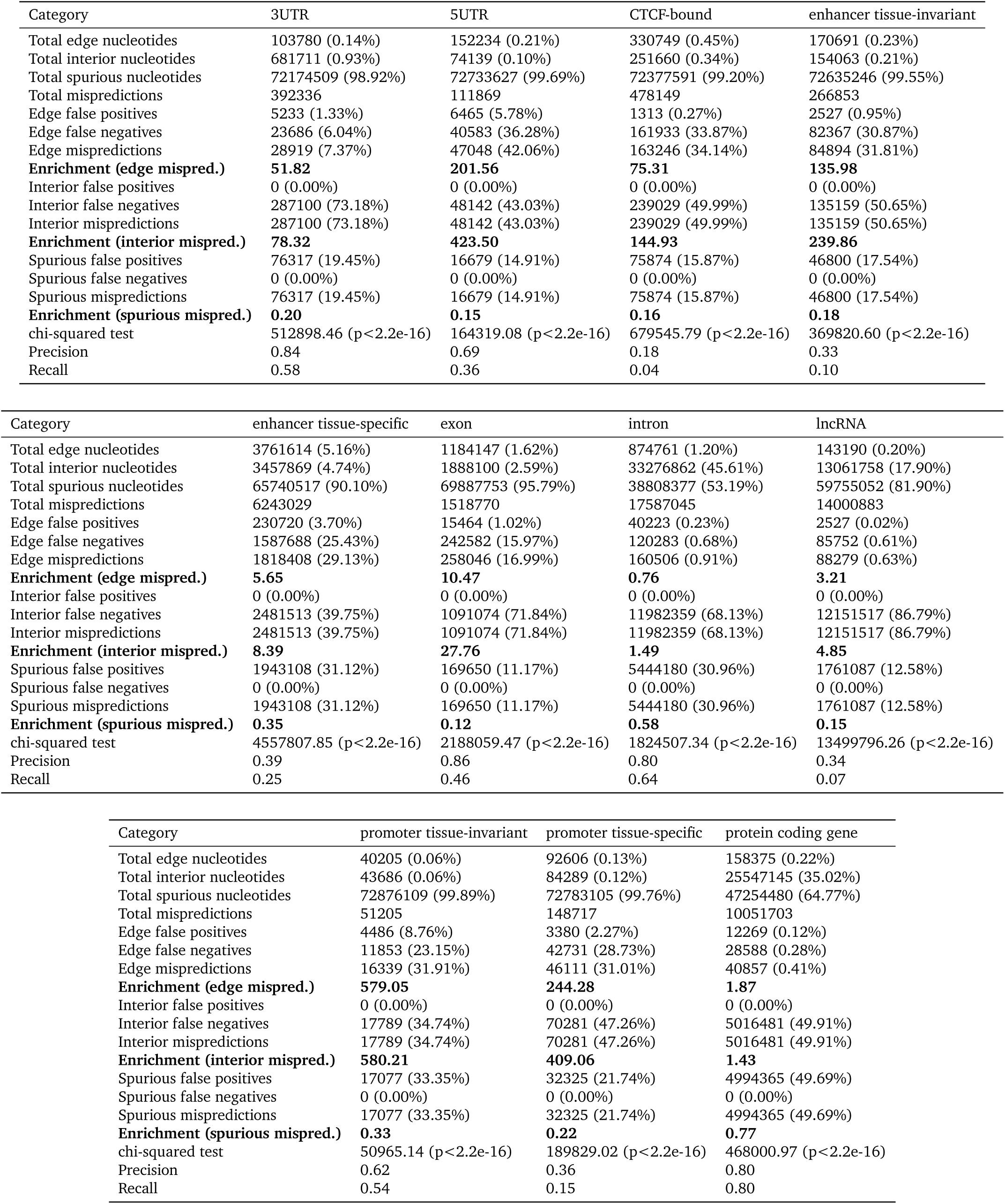
Statistics of types of mispredicted nucleotides by the model SegmentNT-30kb. For each element type, we splatted all nucleotides from the test set according to their distance to the boundaries of the ground-truth regions: a nucleotide is considered part of the edge if closer than 50nt from a region boundary (edge), part of the interior of a region if further way but inside a positive region (interior), and part of spurious regions otherwise. We include the chi-squared test statistics an p-value for the three classes based on their expected frequencies. Splice sites and polyA sites were not inclused as they are too short to define edges.

## Supplementary Figures

**Supplementary Figure 1.**
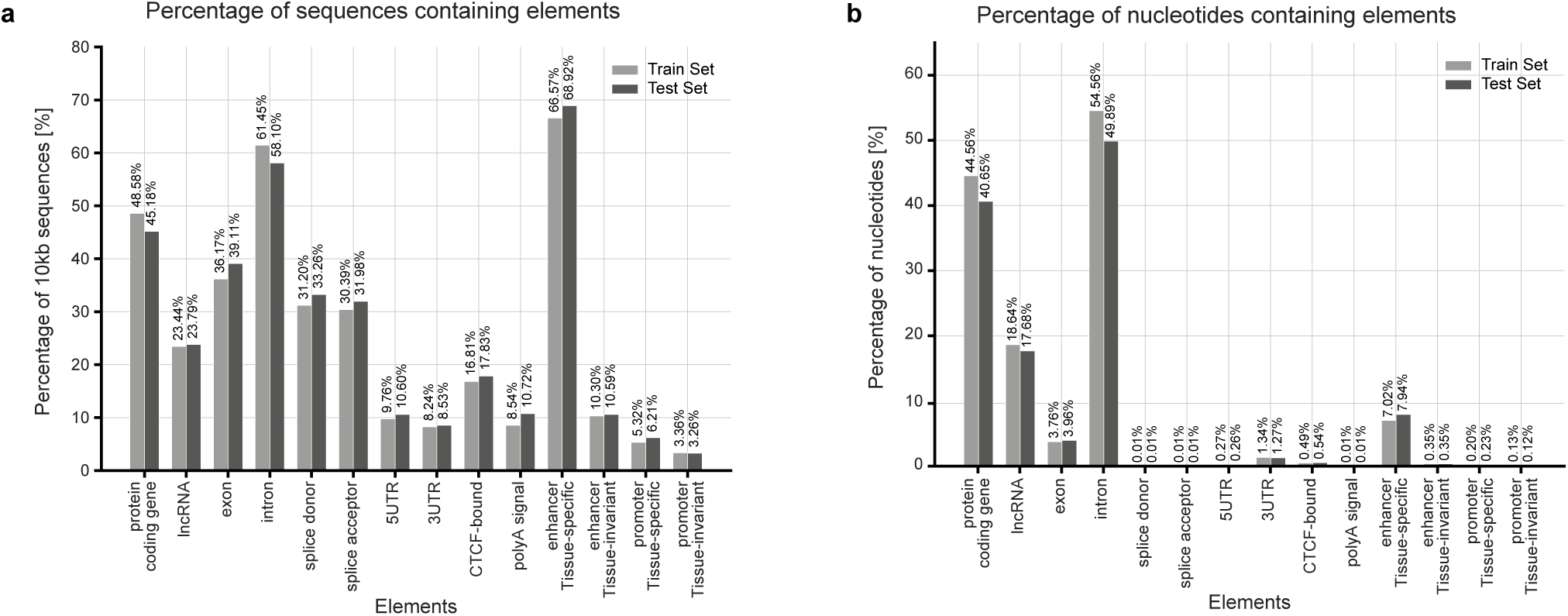
Data distribution per element type. **a)** Percentage of sequences containing each element type in train and test 10kb dataset. **b)** Percentage of nucleotides containing each element type in train and test 10kb dataset.

**Supplementary Figure 2.**
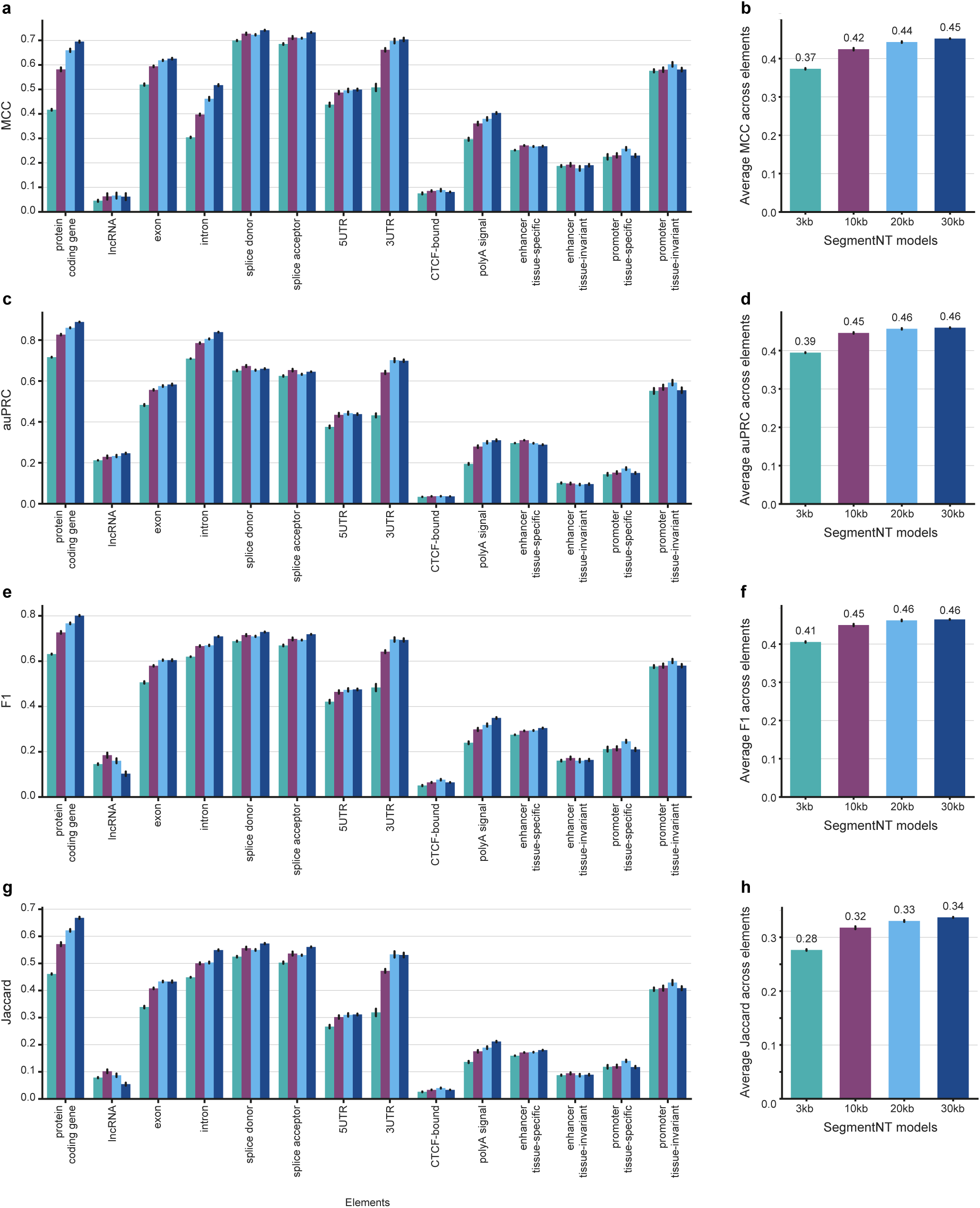
Performance of the different SegmentNT models for different metrics. **a,c,e,g)** Performance of SegmentNT trained on 3kb, 10kb, 20kb and 30kb sequences on 14 types of genomic elements. We used as metrics the MCC **(a)**, auPRC **(c)**, F1-score **(e)** and Jaccard-index **(g)**. Data are presented as mean metric values +/- 95% confidence interval from 10 different samplings of the test set. **b,d,f,h)** Average performance of the different models across the 14 elements. Data are presented as mean metric values +/- 95% confidence interval from the 14 elements.

**Supplementary Figure 3.**
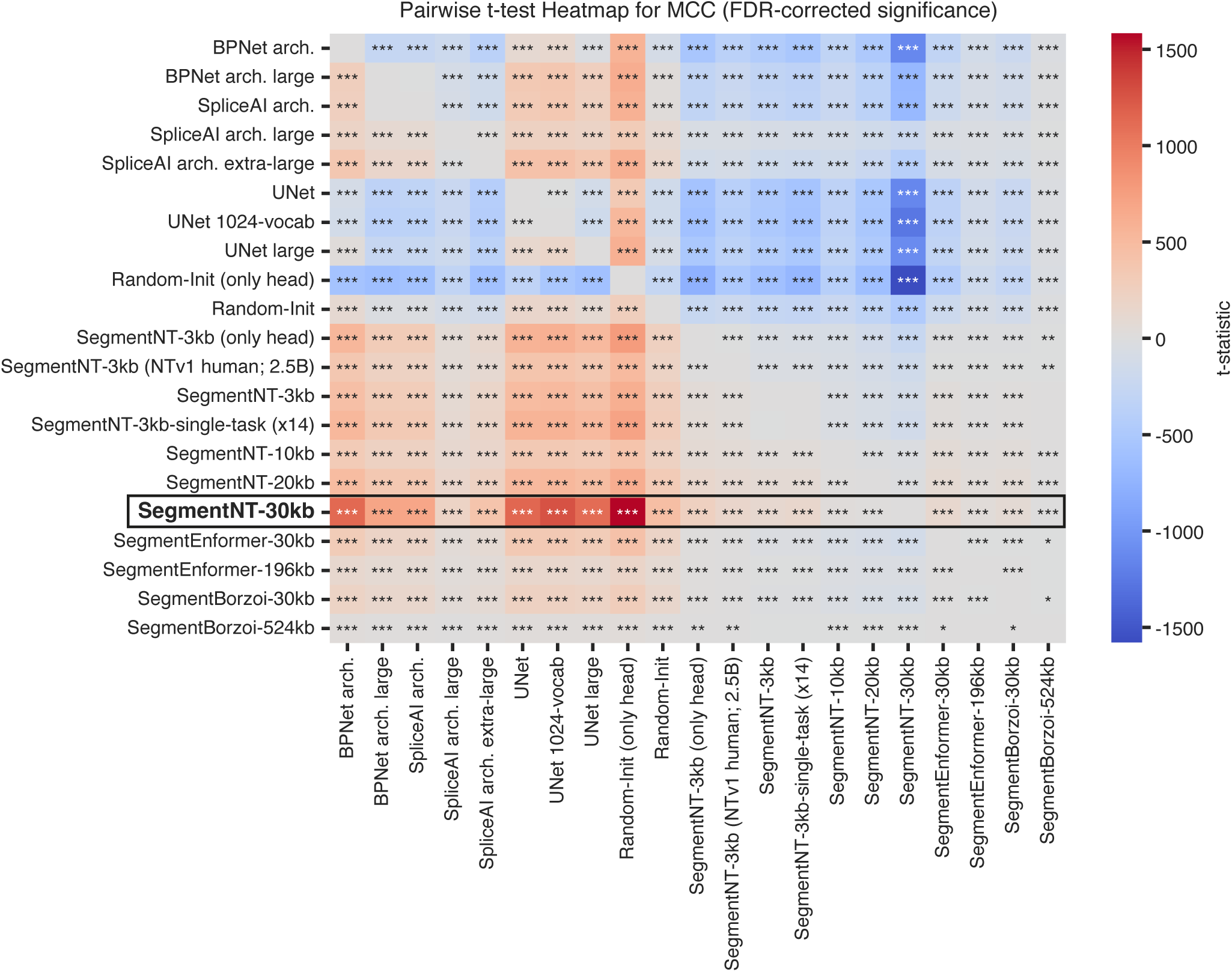
Statistical comparison between all segmentation models. Heatmap displaying the results of pairwise t-tests between different models, comparing the average MCC performance across the 14 features measured over the 10 different data splits. Each cell in the matrix represents the t-statistic, color-coded according to its magnitude and direction. Positive values (red) indicate a higher average in the y-axis label of the comparison, while negative values indicate a higher average in the x-axis label. To control for multiple comparisons, we applied the Benjamini-Hochberg False Discovery Rate (FDR) correction to the resulting p-values. Cells are annotated with significance stars indicating the adjusted p-values: *** for *p* < 0.001, ** *p* < 0.01, * *p* < 0.05, and no stars indicate *p ≥* 0.05.

**Supplementary Figure 4.**
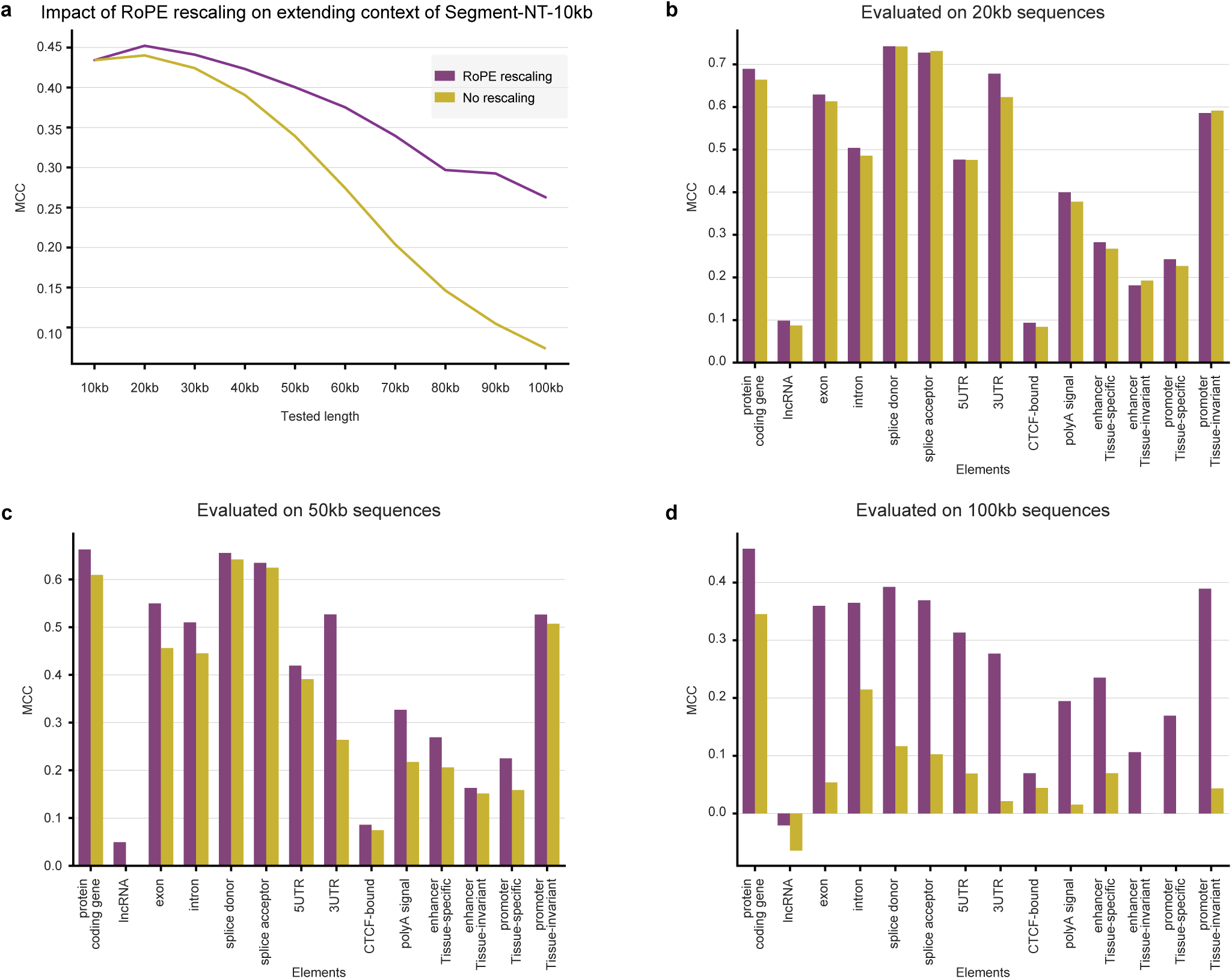
Context-length extension allows to rescale SegmentNT-10kb to 100kb sequences. **a)** Average MCC performance across the 14 elements for the SegmentNT-10kb model with and without context-length rescaling. **b-d)** Performance on different input lengths without vs with context-length extension.

**Supplementary Figure 5.**
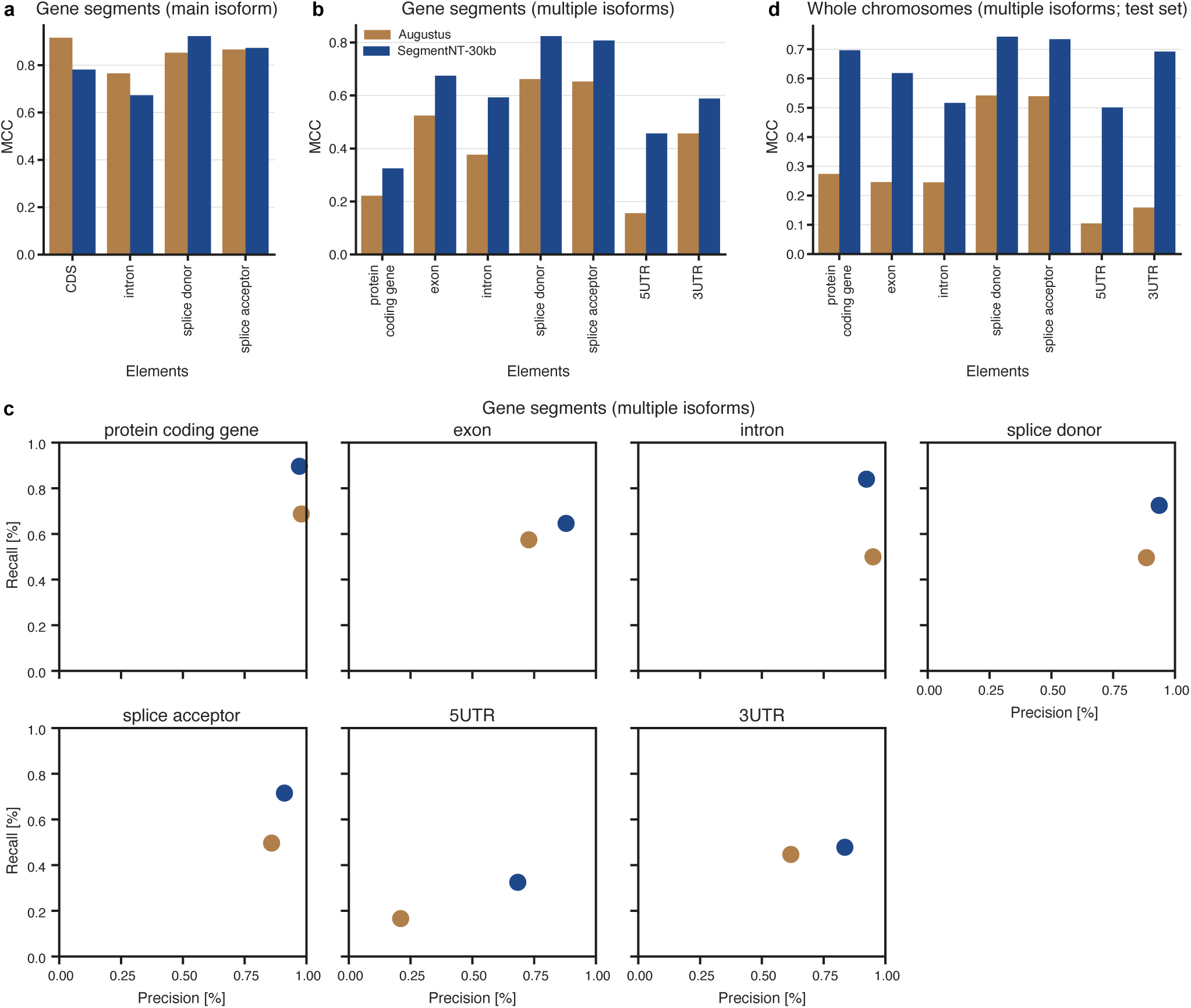
Benchmarking of gene prediction with AUGUSTUS. **a,b,d)** Performance of SegmentNT-30kb and AUGUSTUS for the different gene elements in different datasets: 30kb segments with genes using only their main isoform **(a)**, 30kb segments with genes considering all annotated gene isoforms **(b)** and whole segmentation test chromosomes considering all annotated gene isoforms **(d)**. The metric used was the MCC. **c)** Precision and recall for SegmentNT-30kb and AUGUSTUS for the different gene elements in the dataset of 30kb segments with genes considering all annotated gene isoforms.

**Supplementary Figure 6.**
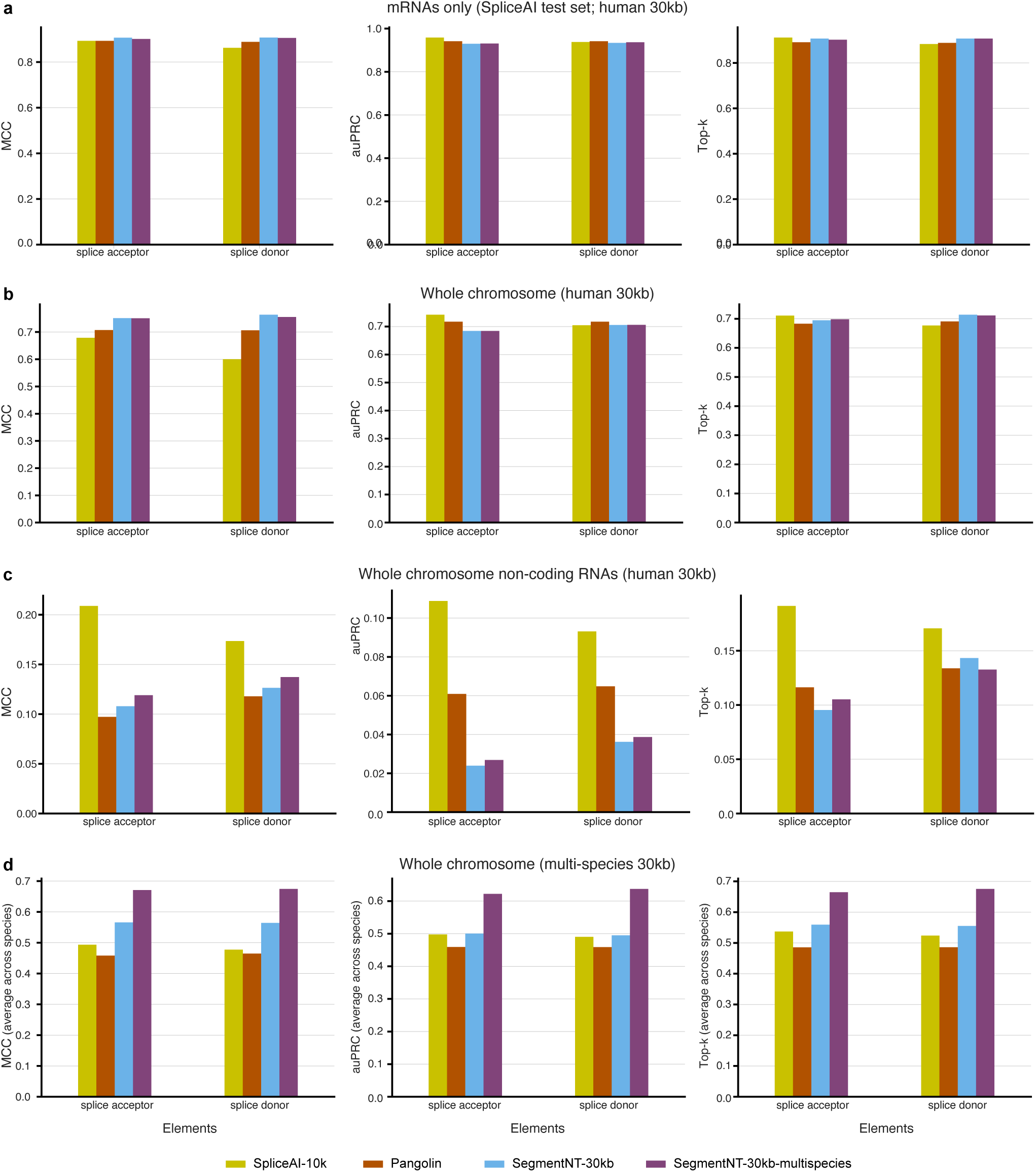
Comparison with different splicing prediction methods. Performance of the SpliceAI, Pangolin and SegmentNT-30kb and -multispecies models for splice acceptor and donor detection. We show MCC, auPRC and Top-k metrics on **(a)** the SpliceAI’s mRNA-based test set, **(b)** the human SegmentNT’s whole genome test sets, **(c)** the same but after removing protein-coding genes to focus on non-coding RNA splice sites; and **(d)** the multispecies SegmentNT’s whole genome test sets. The performance in the multispecies dataset is based on the average across 20 species.

**Supplementary Figure 7.**
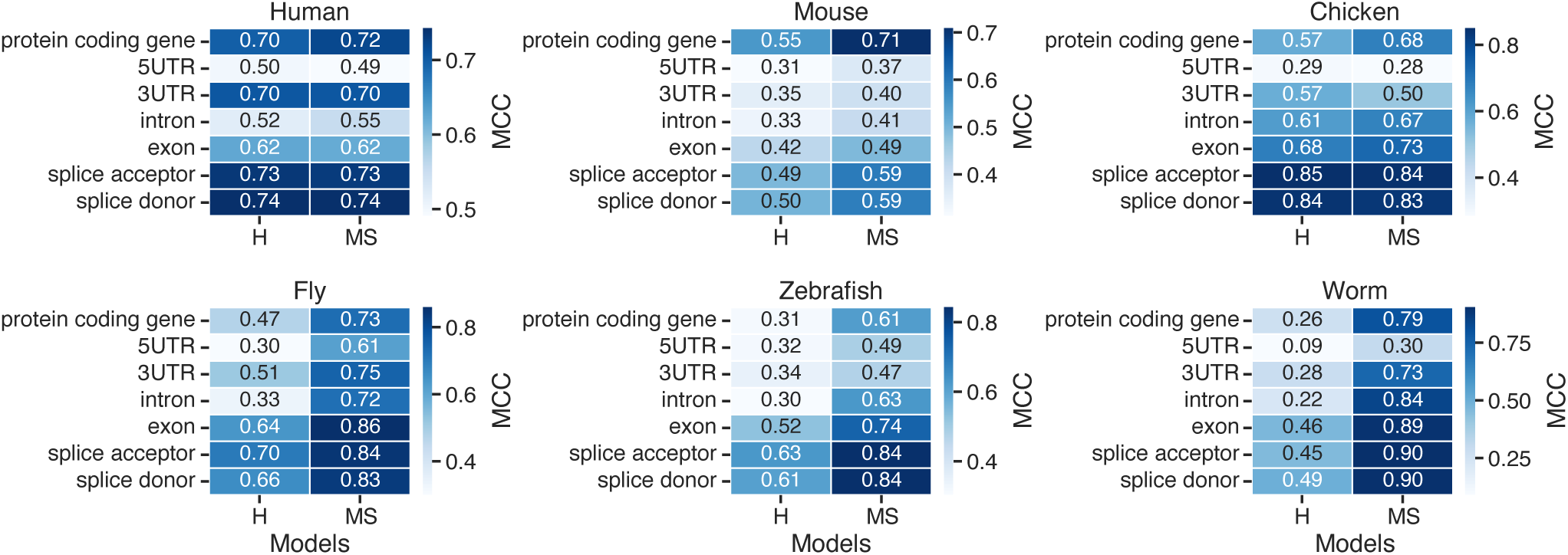
Comparison of human (H) and multispecies (MS) SegmentNT models on training set species. Data is shown as the mean MCC values from 10 different samplings of each species’ test set.

**Supplementary Figure 8.**
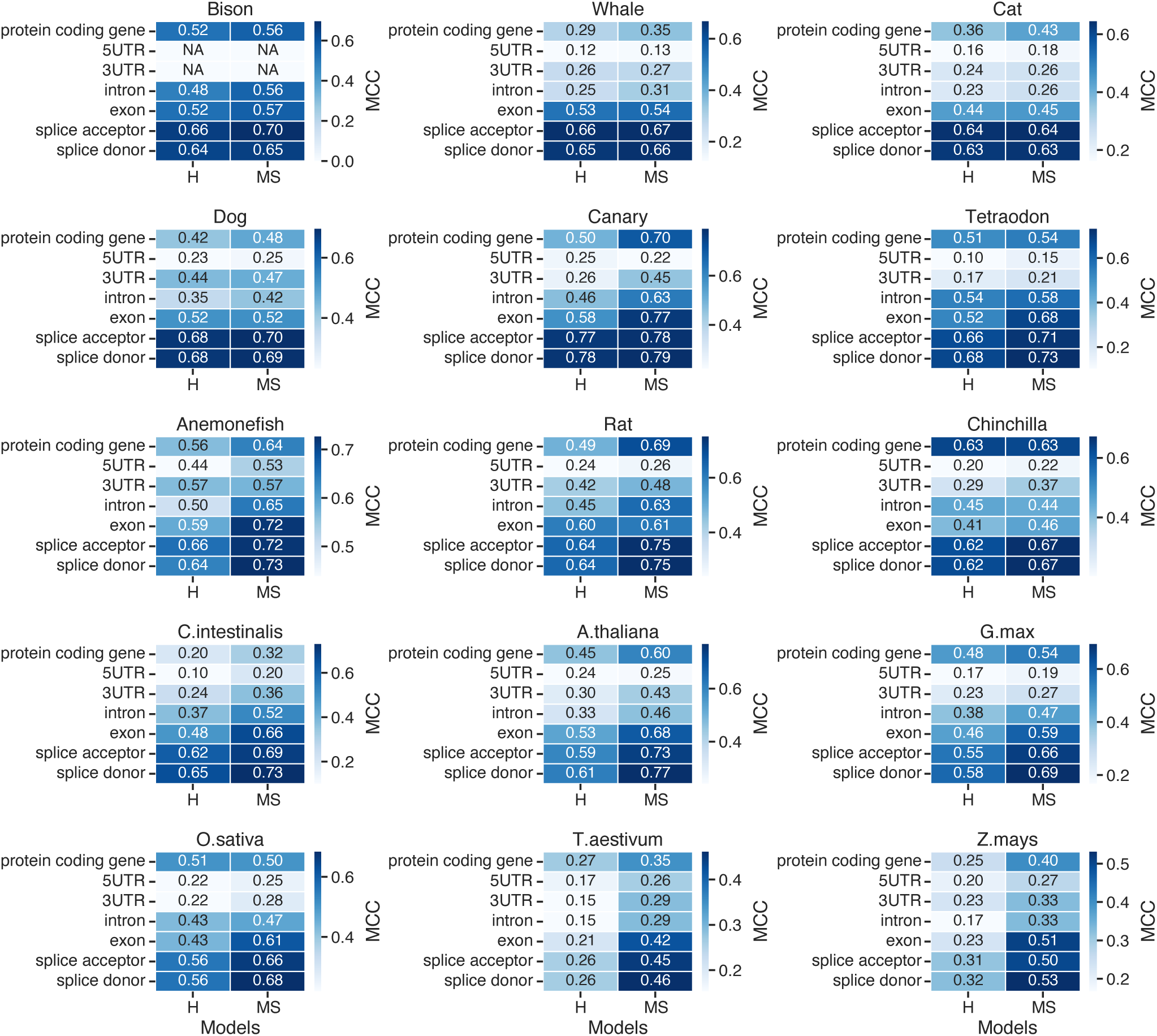
Comparison of human and multispecies SegmentNT models on test set species. Performance of the human (H) and multispecies (MS) model per element for animal and plant test set species. Data is shown as the mean MCC values from 10 different samplings of each species’ test set.

**Supplementary Figure 9.**
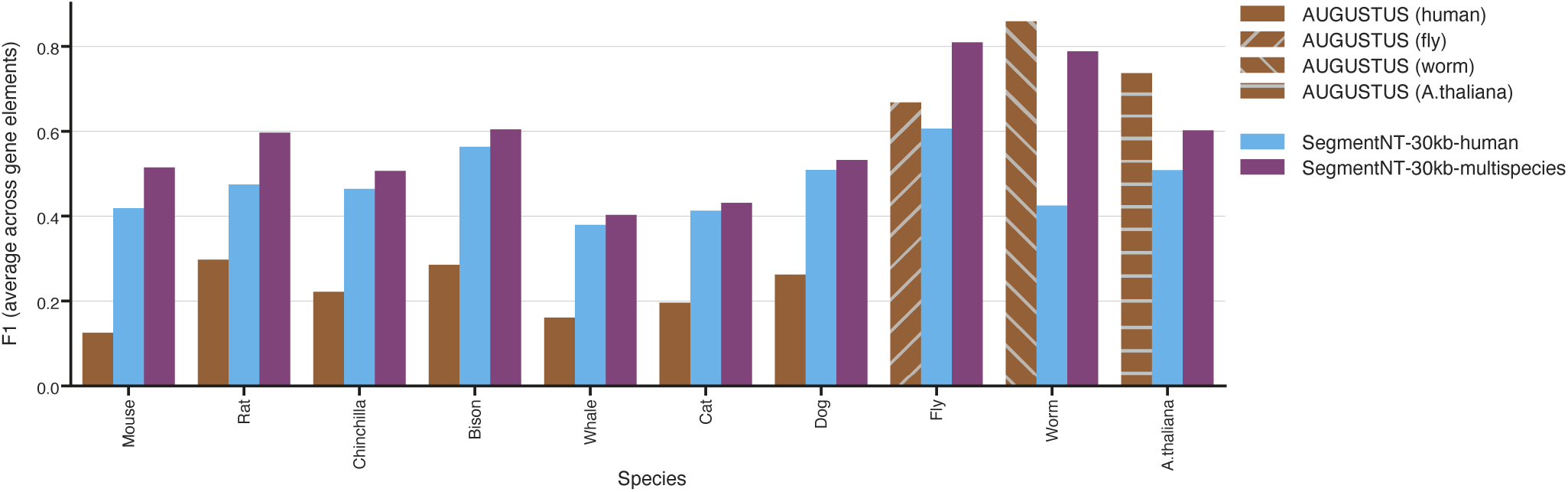
Comparison with AUGUSTUS on gene annotation across species. Performance of AUGUSTUS compared with SegmentNT-30kb human and multispecies models for the annotation of genes in the genomes of the different species. We show the species for which there are AUGUSTUS models: as recommended by AUGUSTUS, we used the human model for all mammals and specialized models for the other species (fly, worm and Arabidopsis). The metric used is the average F1-score across gene elements per species.

## Footnotes

1 https://basespace.illumina.com/projects/66029966/

## Notes

### Summary of Updates

We have fixed a typo in the abstract.

